# Artificially cultivated duckweed: a high-efficiency starch producer

**DOI:** 10.1101/2023.03.23.533730

**Authors:** Yang Fang, Ling Guo, Songhu Wang, Yao Xiao, Yanqiang Ding, Yanling Jin, Xueping Tian, Anping Du, Zhihua Liao, Kaize He, Shuang Chen, Yonggui Zhao, Li Tan, Zhuolin Yi, Yuqing Che, Lanchai Chen, Jinmeng Li, Leyi Zhao, Peng Zhang, Zhengbiao Gu, Fangyuan Zhang, Yan Hong, Qing Zhang, Hai Zhao

## Abstract

The increasing demand for starch has been a social struggle. We report a new technology that efficiently produces starch from duckweed. Although *Landoltia punctata* has a dramatic contraction of gene families, its starch content and productivity reached 72.2% (dry basis) and 10.4 g m^-2^ d^-1^ in 10 days, equivalent to a yield of 38.0 t ha^-1^ y^-1^ under nutrient limitation and CO_2_ elevation treatment. Meanwhile, we also examined the mechanism of duckweed’s high starch accumulation. This is exhibited in the regulation of DNA methylation and transcription factors as well as the significantly up-regulated transcription levels and the increased enzyme activities of key genes in starch biosynthesis. Meantime, while nitrogen redistribution was enhanced, sucrose biosynthesis and transportation, and lignocellulose biosynthesis were all reduced. These alterations led to a reduction in lignocellulose and protein content and ultimately an increase in an accumulation of starch in the chloroplast. This work demonstrates duckweed’s potential of being a highly efficient starch producer.

## 1 Introduction

Starch plays a pivotal role in the human society. It provides 80% of the world’s calories ^1-3^ while serving as raw materials for biochemical industry and biofuel production ^4^. Starch primarily originates from starch-storing organs of staple crops ^1^ and accounts for 60-70% of these storage ^5^. Thus starch productivity is highly correlated to the crop yields ^2^.

The estimated annual yield of staple crops is 2.5 billion tons worldwide ^6^, mainly including three types of starch-storing organs: cereal grains (e.g. corn, wheat, rice, and barley), root and tubers (e.g., potato, sweet potato, yam, and cassava), and beans ^6^. However, these organs only comprise of parts of the whole crop plants ^7^, whereas other parts, such as stems and leaves, are agricultural residues and wastes that may become environmental pollutants. Cereal grains, as the most important part of crops, are the main sources of starch. As seed organs, cereal grains’ development depends on sexual reproductive growth. This process, including flowering, pollination, and grain-filling, can be easily disrupted by biotic and abiotic stresses ^8^.Therefore, stable production of cereals and beans has beena challenge. Further, although the starch biosynthetic pathway has been well studied, genetic manipulations to significantly increase starch productivity remain difficult due to the complexities of starch metabolic networks ^2,6^. With the continuous growth of the world’s population, the demand for staple crops is predicted to rise by 70-100% by 2050 ^9,10^. The sustainable staple crop supply has been and will always be a challenge. Therefore, it is urgent to search for new approaches e for high-efficient starch production with new starch crops.

Duckweed, a floating aquatic monocot, is the fastest-growing higher plant on the earth ^11^. Without stems, it consists of “frond” structures and a few or no roots. It accumulates biomass through asexual budding and vegetative growth processes ^11-13^. Its biomass can increase near-exponentially and their estimated yield reaches 55 t ha^-1^ per year (dry weight, DW) ^14^. Duckweeds are a feed source for domestic animals, fishes, and even food source for indigenous people in Southeast Asia ^15^. Duckweeds have caught extensive attentions because of their potential application in feed/food, bio-energy production, and wastewater treatment ^16^. It was reported that D Researches show that duckweed can reach 75% starch content in a sugar substrate ^17,18^. Especially under the cultural conditions without any organic carbon, it can reach 48% starch content after 10-day treatment (sugar-free solution) ^17^. These findings indicated its potential as a new starch crop for biofuel conversion and food supplement. It is necessary to further improve the starch production capacity, elucidate the mechanism of its starch accumulation, and evaluate the potential of duckweed.

Herein we develop a novel duckweed cultivation technology that requires limited nutrients and CO_2_ supplementation to achieve very high starch content (>72.2%) and extremely efficient production. In this study, we integrated whole-genome sequencing, epigenomics, transcriptomics, enzyme activity, and composition variation. With these methods, we uncovered the mechanisms of efficient starch accumulation in duckweed in starch accumulation and carbon partitioning, gene expression regulation of starch metabolic pathway, and sucrose biosynthesis and transportation.

## 2 Results and discussion

### 2.1 A simple technology for efficiently producing starch in duckweed

We developed a simple technology that makes duckweed an efficient starch-producer. *Landoltia punctata0202*was previously identified to be a useful duckweed ecotype with high potential for starch accumulation ^17,19,20^. We used limited nutrients (“L” condition by cultivating duckweed in deionized water) and an elevated concentration of CO_2_ (“C” condition, by supplying CO_2_ to 2500±100 ppm) to stimulate biomass accumulation and starch production (Appl. No. ZL201710855019.8). This technology greatly improves the starch content, enhances the biomass accumulation, and dramatically increases the starch yield in duckweed. The starch content increased from 6.5% to 72.2% (dry basis, d.b.) (Fig. 1A). The biomass of duckweeds reached

**Fig. 1.**
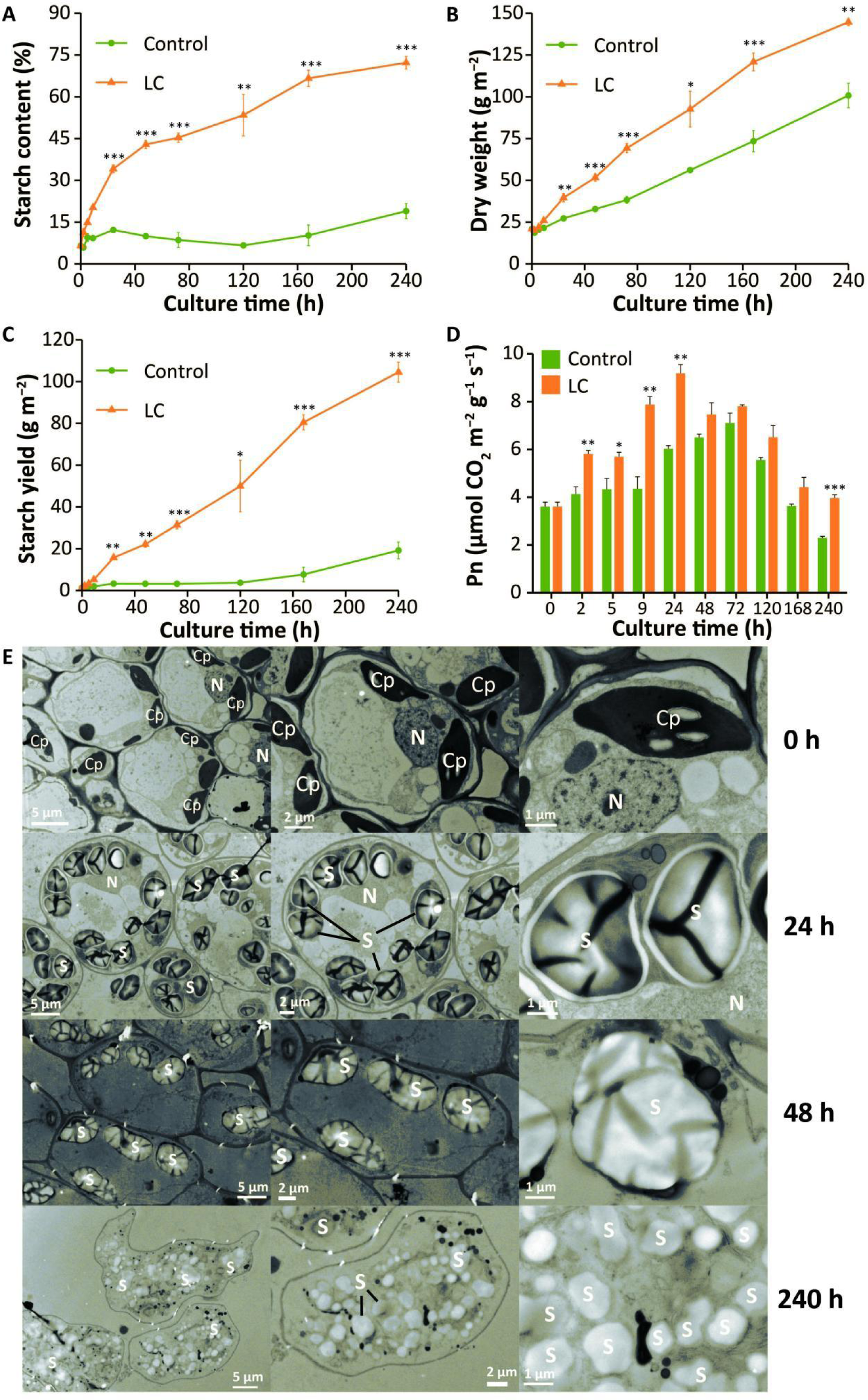
Accumulation of starch in *Landoltia punctata* under LC treatment. **A-D,** Changes of (A) starch content, (B) dry weight, (C) starch yield and (D) net photosynthetic rate during 240-hour cultivation. Control, cultivated in 1/5 Hoagland medium. LC, cultivated under conditions of nutrient limitation and elevated CO_2_ (2500±100 ppm). Pn, net photosynthetic rate. Error bars represent the standard deviation measured from three independent cultures. Asterisks indicate represents statistically significant difference comparing data from each treatment group with control in the same assay conditions (Student’s *t*-test). *, P<0.05; **, P<0.01; ***, P<0.001. **E**, Starch granules in duckweed fronds observed by transmission electron microscope (TEM). Duckweed was cultivated under LC treatment. The fronds were sampled at 0, 24, 48 and 240 h, then fixed, embedded, and dehydrated prior to observing by TEM. Cp, chloroplast; S, starch; N, nucleus.

144.7 g m^-2^ (DW) in 10 days, and that of control was only 100.8 g m^-2^ (Fig. 1B).A net amount of 104 g starch was produced per square meter (Fig. 1C), equivalent to 38.0 t ha^-1^ y^-1^, higher than that of almost all storage organs of crop plants ^21,22^. There have been no reports that nutrient limitation can simultaneously increase the starch content and yield in cereals. Studies on the model plants also indicated that, although nutrient limitation can increase the starch content, it remarkably reduced the plant biomass at the same time ^23^.

During the cultivation process, the fronds of duckweed gradually became larger and distinctively turned yellow under limited nutrients and CO_2_ supplementation (LC) treatment (Fig. 2D) Its moisture content gradually decreased - from 90.98±0.52% at the beginning to 69.33±0.71% at the 10th day (Fig. S2).

**Fig. 2.**
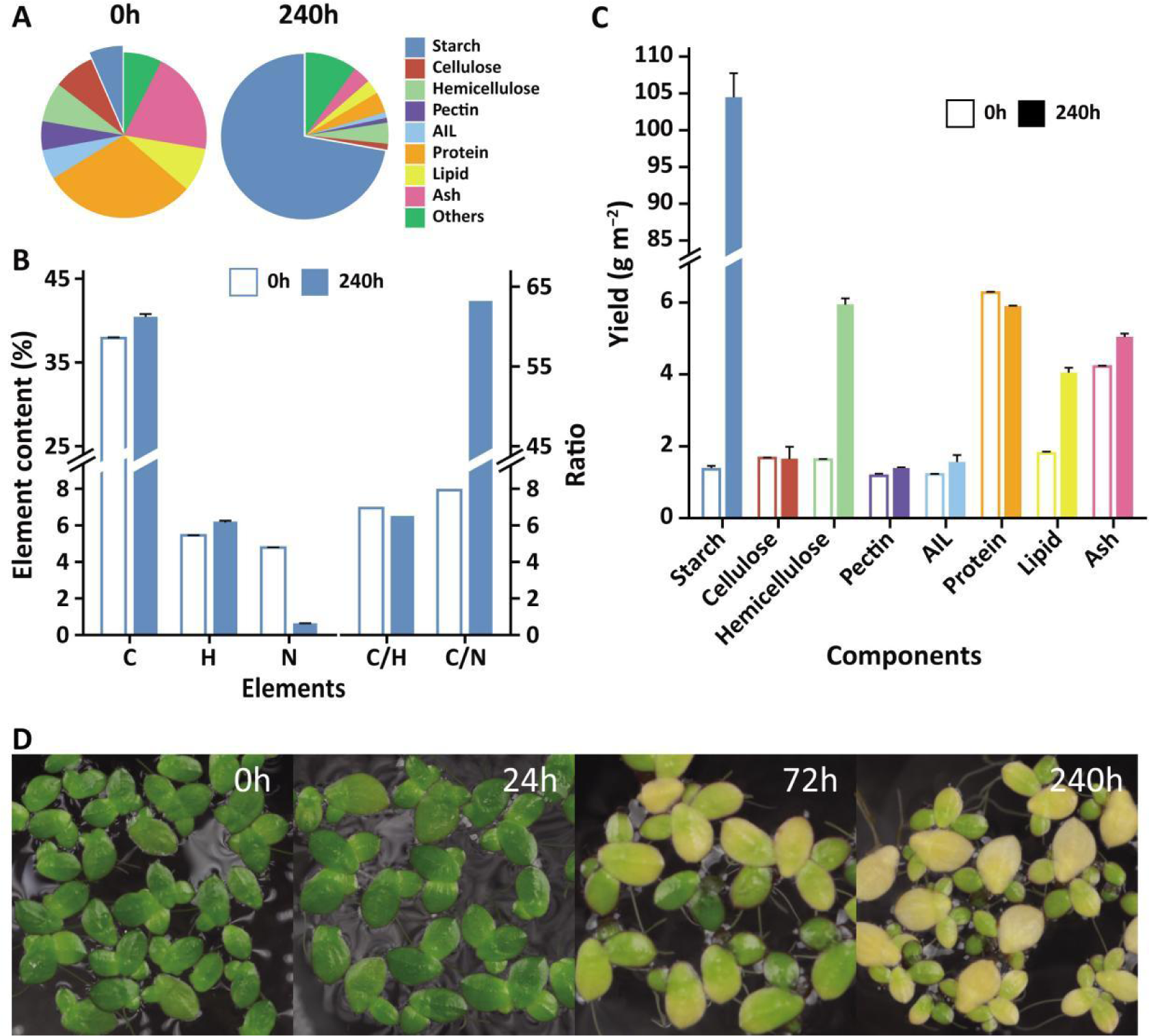
Changes in content and yield of primary composition in *Landoltia punctata* under LC treatment. **A,** The content changes of primary compounds in duckweed before treatment and 240 hours after treatment. AIL, acid insoluble lignin. Lipid was extracted using diethyl ether. **B,** Changes of primary elements content and their ratio in duckweed before treatment and 240 hours after treatment. C, carbon; H, hydrogen; N, nitrogen; C/H, the ratio of carbon to hydrogen content; C/N, the ratio of carbon to nitrogen content. **C**, The yield changes of primary compounds in duckweed before treatment and 240 hours after treatment. Error bars represent the standard deviation measured from three independent cultures. **D**, Fresh fronds at different culture time under treatment.

Meanwhile, the net photosynthetic rate (Pn) of duckweed increased at the beginning and then gradually decreased, and the Pn of the treatment group was always higher than that of the control group within the 10-day period. (Fig. 1D).

### 2.2 Starch accumulation and carbon partitioning

#### 2.2.1 Expression and activities of key enzymes in the starch biosynthetic pathway

The transcript levels of all the key starch biosynthesis genes in chloroplast were up-regulated under LC treatment (Fig. 3A). Plastidial ADP-glucose pyrophosphorylase (AGPase) (Fig. S6), as the first rate-limiting enzyme in starch biosynthesis, determines carbon flux into starch to a large extent. AGPase is precisely regulated at transcriptional and post-translational levels including allosteric regulation and redox modulation ^24,25^. Previous studies revealed that the activity of AGPase is induced by increased 3-phosphoglycerate (3-PGA) and decreased phosphate through allosteric regulation ^26,27^ and suppressed by phosphate ^28^ and nitrate ^29^ through transcriptional regulation. In our case, elevated CO_2_ significantly increased 3-PGA content from 162.4 μg g^-1^ fresh weight (FW) to 292.2 μg g^-1^ FW (Fig. S10A) The nutrient limitation caused the deficiency of phosphate and nitrate. The combination of the activation (mediated by increased 3-PGA and reduced phosphate) and the inhibition (mediated by the release of phosphate and nitrate to the improvement of AGPase gene expression and enzyme activity (Fig. 3B, Fig. S10B, and Fig. 3E). AGPase had 3.0× expression and 6.5× activity under LC treatment (Data S2, Fig. 3A and B). Most previous attempts on increasing starch accumulation only focused on enhancing AGPase gene expression instead of regulating its enzyme activity. This perhaps is the reason why they were difficult to succeed ^30-32^.

**Fig. 3.**
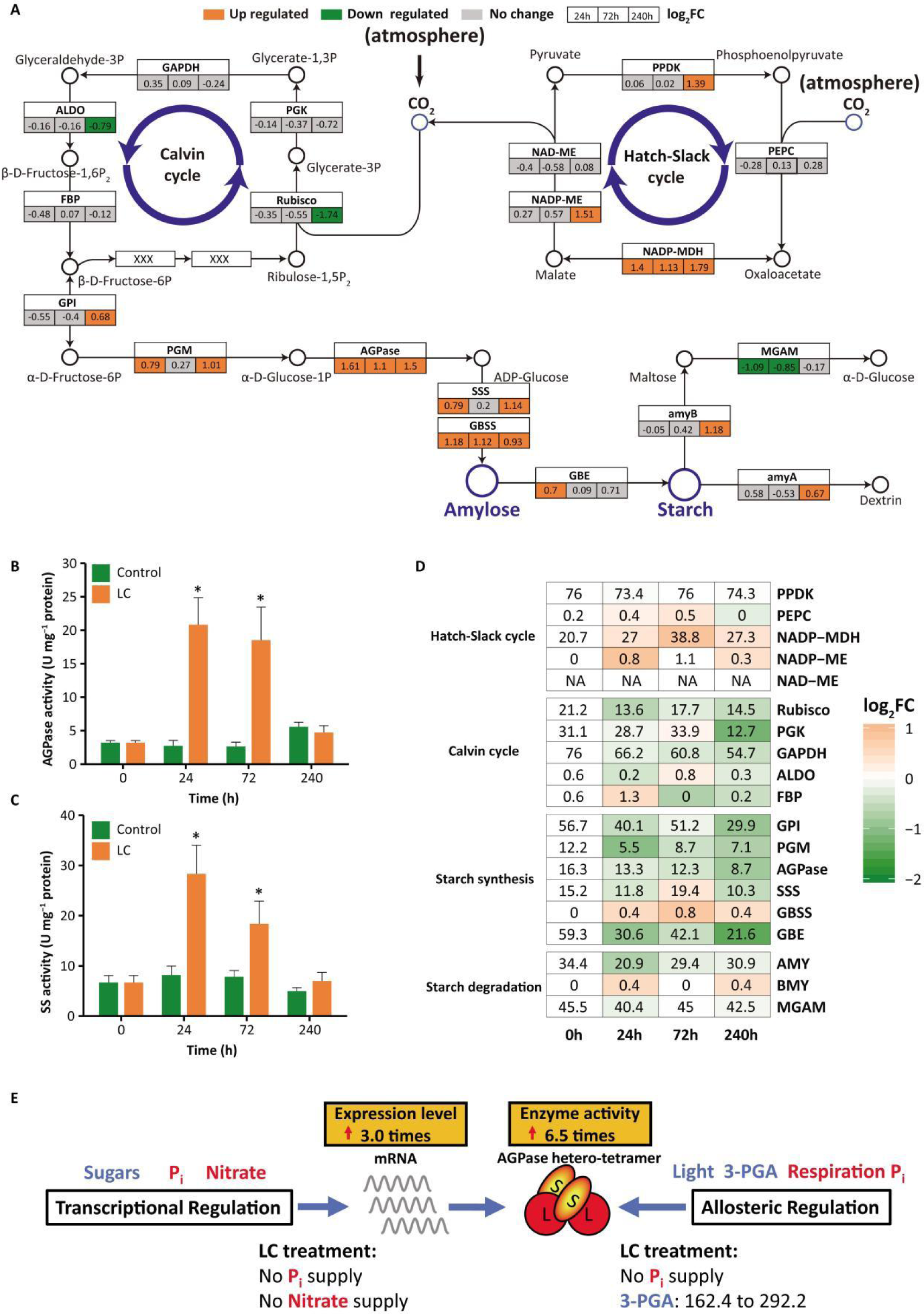
Expression and DNA methylation of genes involved in starch metabolism in *Landoltia punctata* under LC treatment. **A,** Expression of key genes involved in CO_2_ fixation, carbon concentration, starch biosynthesis, and starch degradation. Boxes colored in orange or cyan indicated the up-regulated or down-regulated differentially expressed genes (p value ≤ 0.05, |Log_2_(fold change)| ≥ 0.58) after cultivated for 24 h, 72 h, and 240 h, as compared with that at 0 h. Numbers presented in the box show the log_2_FC values. **B and C,** Changes of key enzyme activities in starch biosynthesis pathway at different culture time under treatment. AGPase, ADP-glucose pyrophosphorylase; SS, starch synthase, including both SSS (soluble starch synthase) and GBSS (granule-bound starch synthase). Error bars represent the standard deviation measured from three independent cultures. Asterisks indicate significance of difference as compared with control (evaluated by one-way analysis of variance, ANOVA). The unlabeled data indicates insignificance. *, P<0.05. **D,** DNA methylation rates in 2 kb-upstream region of key genes in CG context involved in CO_2_ fixation, carbon concentration, starch biosynthesis, and starch degradation. Numbers in the box indicated the methylation rates (%). The color gradient indicates log_2_FC of methylation rate comparing with 0 h, where FC is fold change. **E,** Schematic diagram of AGPase activity regulation in duckweed under treatment. AGPase activity is regulated in transcriptional and allosteric level through multiple environmental factors (3-PGA, Pi, and nitrate) under LC treatment. Blue font and arrow indicate activation, red font indicates inhibition and red arrows indicate the increase of expression or enzyme activity. 3-PGA, μg g^-1^ FW.

Furthermore, LC treatment also increased the expression and activities of other important genes/enzymes including granule-bound starch synthase (*GBSS*), soluble starch synthase (*SSS*), and also the expression of 1,4-alpha-glucan branching enzyme (*GBE*) (Fig. 3A and C). Previous studies showed that, the genetic manipulation of any one of the genes mentioned above usually had limited effects on starch content and no increase in net starch yield ^6,33-36^. Starch biosynthesis is a complex system that interconnects with a wide variety of cellular processes and metabolic pathways ^24^. Thus, it is very difficult to develop a comprehensive method to regulate it. In our case, the biomass increased by 6x, the starch content increased by 11xand the starch yield increased by 76xwithin 10 days compared to the beginning stage (Fig. 1 and 2). This is presumably because LC treatment could significantly improve the activities of AGPase by transcriptional regulation and allosteric regulation, increase the gene expression level and activity of other key enzymes, and finally achieve the regulation of the complex system.

#### 2.2.2 Distribution of carbon on lignocellulose and protein

The carbon skeletons of lignocellulose and protein derive from photoassimilates, and their biosynthesis is strongly affected by carbon partitioning. Lignocellulose contents in duckweeds are relatively low and were further reduced under LC treatment. (Fig. 2A). Initially, cellulose, hemicellulose, and pectin contents were 8.0±0.1%, 7.8±0.1%, and 5.6±0.3% (d.b.) After ten days of treatment, they decreased to 1.1±0.2%, 4.1±0.1%, and 1.0±0.0% (d.b.) (Fig. 2A), a reduction of 86.3%, 47.4%, and 82.1%, respectively. The expression level of sucrose synthase (SUSY), an enzyme involved in the formation of UDP-glucose from sucrose to cellulose and hemicellulose, was significantly down-regulated (Fig. S12 and Data S7). Regarding cellulose degradation, the expression of genes in glycoside hydrolase family 9 (cellulase) was significantly up-regulated to more than 5x(Data S8). These changes might contribute to the reduction in cellulose and hemicellulose content.

Lignin, a complex phenol polymer hard to be degraded, is the main obstacle for biomass utilization. Lignin content in duckweed decreased 81.0% (Fig. 2A) from 5.8±0.1 to 1.1±0.1% under 10 days of LC treatment. The gene expression of laccase, the key enzyme for monolignol polymerization and crosslinking, was extremely low (FPKM values < 20) (Fig. S12 and Data S9), providing a possible explanation for the low lignin content. Also, lignocellulose composition was reduced to a very low level (6.3%), indicating a higher quality of whole biomass (Fig. 2A).

Notably, the protein content of duckweed dropped rapidly from 30% to 4% (d.b.) under this treatment (Fig. 2A). The total amount of protein remained basically stable within 10 days due to the lack of nitrogen supply (Fig. 2C) while the biomass, especially starch content, of duckweed still accumulated rapidly (Fig. 1B and Fig. 1C). Nitrogen glutamine synthetase (*GS*), the key gene in nitrogen assimilation and recycling, plays an important role in increasing nitrogen use efficiency (NUE, kg grain yield per kg N application) of crops and increasing cereals yield ^37,38^. The expression level of *GS*was up-regulated by 6.7xand its enzyme activity was 7.4x higher than that of the control (Data S12 and Fig. S11). Previous studies proved that nitrogen limitation significantly reduces mRNA expression of *GS* and its enzyme activity and increases starch content without increasing the plant biomass and the starch yield ^23,39^. However, in this study, the plant biomass and starch yield both increased possibly by increasing the expression of *GS*, which greatly promoted the redistribution of ammonium and therefore enhanced the protein reuse (Fig. S13). Thus, future research could focus on duckweed *GS*, especially on its role in improving the NUE.

Reduction of protein and lignocellulose content in duckweeds matched well with the down-regulated expression of relevant genes. This and the increase of starch content indicated that a large amount of photoassimilate flowed to starch synthesis under LC treatment. Therefore, duckweed offers an ideal model for nitrogen and carbon metabolism regulation study.

### 2.3 Gene numbers and regulation of carbon assimilation pathways

#### 2.3.1 Contraction of total gene numbers in carbon assimilation

We analyzed duckweed’s genes of the carbon assimilation pathway because of its strong starch accumulation ability. According to the results of whole-genome sequencing of *Landoltia punctata*0202 (GeneBank accession number: PRJNA546087) and the corresponding gene family analysis (Data S1), the total gene number of carbon assimilation is only 50, including starch metabolism, Calvin cycle, and Hatch-Slack cycle. This is significantly contracted compared to those in Arabidopsis (64), rice (82), and maize (86) (Table 1). Under LC treatment in the carbon assimilation pathways, only in starch metabolism were the transcript levels of the key starch biosynthesis genes and the corresponding enzyme activities enzyall up-regulated. (Fig. 3A-C and Fig. S3). These starch biosynthesis genes worked in a synergistically efficient way, hence enhancing starch formation. There was almost no change in the gene expression in the Calvin cycle and also no significant change in the activity of its key enzyme Rubisco (Fig. 3A and Fig. S8C). In Hatch-Slack cycle, part of the genes such as *NADP-MDH*, *NADP-ME,*and *PEPC*were also up-regulated (Fig. 3A), with the same tendency shown in the corresponding enzyme activities (Fig. S8D-F). LC treatments mainly enhanced the starch biosynthesis, and also had a certain effect on CO_2_ concentration, but had no obvious effect on Calvin Cycle.

**Table 1.**
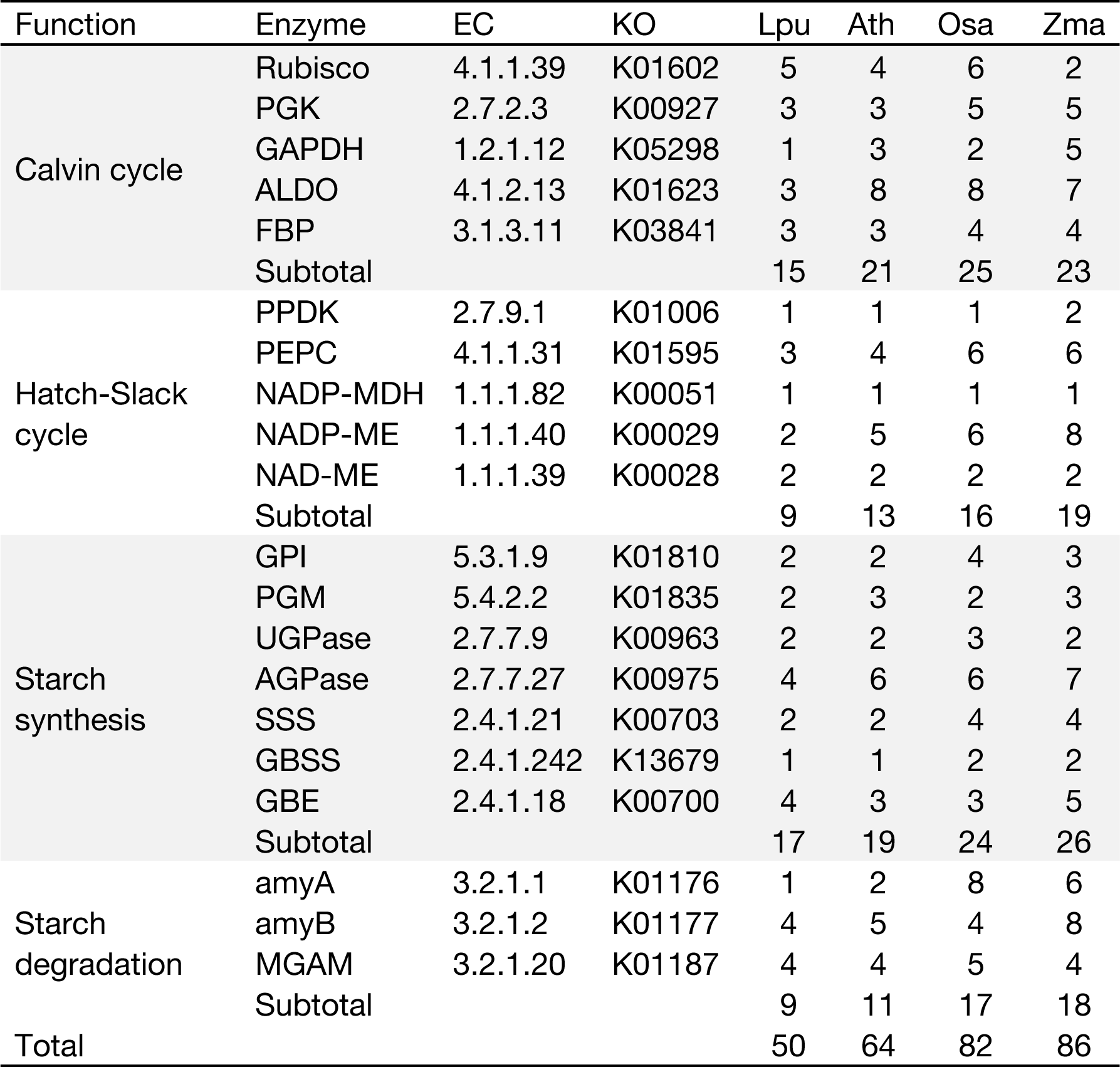
Numbers of genes involved in starch metabolism. Lpu, *Landoltia punctata*; Ath, *Arabidopsis thaliana*; Osa, *Oryza sativa*; Zma, *Zea mays*.

Therefore, what is observed on duckweed contradicts the common knowledge that the more genes an organism possesses, the more function it has. Our results displayed that the changes in starch accumulation may be resulted from the regulation of starch biosynthetic pathway, rather than the number of gene copies.

#### 2.3.2 DNA methylation in gene regulation

DNA methylation, a conserved epigenetic modification, plays an important role in assisting in gene regulation and genome stability ^40^. We studied the epigenome of DNA methylation in the same samples of transcriptome analysis. LC treatments decreased 1.7% of the DNA methylation level in the whole genome of duckweed, from 12.9% to 11.2% (mC) in 24 h (Table S6). More importantly, the DNA methylation level of *AGPase*, *SSS,*and *GBE*promoters were significantly reduced by 46.6, 32.2, and 63.6% (mCG) respectively, while their expressions were significantly up-regulated. This means that in starch biosynthetic pathway, the DNA methylation in the promoter regions experiences a negative correlation with the gene expression (Fig. 3A, D). In the Calvin cycle, although the DNA methylation level of the key genes’ promoters decreased, their expression levels did not change significantly. In the Hatch-Slack cycle, the DNA methylation level of the same key genes’ promoters significantly increased, while their expression levels did not decrease. Notably, the expression level, DNA methylation level, and enzyme activity of NADP-MDH, one of the key genes in Hatch-Slack cycle, all increased significantly (Fig. 3A, D, and Fig. S8E). Previous studies indicated that CO_2_ elevation can significantly up-regulate the activity of NADP-MDH ^41,42^. It could be speculated that the increase in NADP-MDH expression level under LC treatment is due to the effect of elevated CO_2_.

Under LC treatment, DNA methylation did not affect the expressions of key enzymes in the Calvin cycle and Hatch-Slack cycle and only played an essential role in the coordinated expression of the key genes in the starch biosynthetic pathway.

#### 2.3.3 Transcription factors (TFs) in the starch biosynthetic pathway

TFs also play an important role in regulating gene expression ^43^. By analyzing co-expression networks analysis with key genes of the starch biosynthetic pathway, we predicted that some TFs were positively correlated. We found that multiple OBF1, the TFs of the bZIP family in duckweed, were positively correlated with all key genes of starch biosynthesis (*AGPase*, *SSS*, *GBSS*, *GBE*) and they had high expression levels (FPKM increased from 140 to 400) (Data S16). The bZIP TFs are key regulators of starch biosynthesis genes in rice, maize, and wheat and determines starch quality and quantity in endosperms *^44-47^*.

The genes of carbon assimilation are all contracted (Table 1), but under LC treatment, only the starch biosynthesis ability is significantly enhanced, which is mutually confirmed by the changing trends at multiple levels (Fig. 1A, C and Fig.3). This means that high starch yield can be obtained merely by the regulation of expression levels. This discovery deserves further consideration and research.

### 2.4 Sucrose biosynthesis and transportation in duckweed

#### 2.4.1 The “source-flow-sink” relation in plants

Sucrose biosynthesis and transportation is crucial for starch accumulation in plants ^48^. It primarily includes the photoassimilates synthesis in chloroplasts, the transmembrane transport into the cytoplasm, sucrose biosynthesis, and long-distance transportation of sucrose for subsequent conversion into starch in storage organs. The “source-flow-sink” relation is highly related to the crop yield. Our results demonstrated that the duckweed frond, a tissue similar to leaves, acts as a “sink” organ that accumulates storage starch (Fig. 1E).

#### 2.4.2 Conversion of “source” to “sink” in duckweed chloroplast

The transportation of photoassimilates from chloroplast relies on the triose phosphate/phosphate translocator (*TPT*), glucose transporter (*PGT*), and maltose transporter (*MEX*). Correspondingly, deletions and mutations of *TPT*, *PGT*, and *MEX* cause starch accumulation in the chloroplast ^49-51^. Duckweed has a markedly contracted number of genes involved in the transportation of photoassimilates. The number of TPT, PGT, and MEX were reduced to only one copy eachFig. 4 and Data S3). Furthermore, the expression levels of both *TPT* and *PGT* were significantly down-regulated under LC treatment. Notably, *TPT*, the major export transporter of photoassimilates from chloroplasts, decreased by 42.6%(Fig. 4). In duckweed mesophyll cells, LC treatment significantly increased the expression of *AGPase* by 3.0xin the chloroplast while suppressed the exportation of plastidial triose phosphorate and glucose to cytosols, resulting in hyperaccumulation of starch in chloroplast (Fig. 1E, Fig. 3E and Fig. 4). Subcellular localization of starch granules, AGPase, TPT, PGT, and MEX also confirm the transition of chloroplast’s function from being the “source” to the “sink” in duckweed (Fig. 1 and Fig. S6). The source and the sink are now spatially organized together in the chloroplast of duckweed a prominent difference from other crops. This treatment allows duckweed’s chloroplast to be highly efficient in forming and storing starch.

**Fig. 4.**
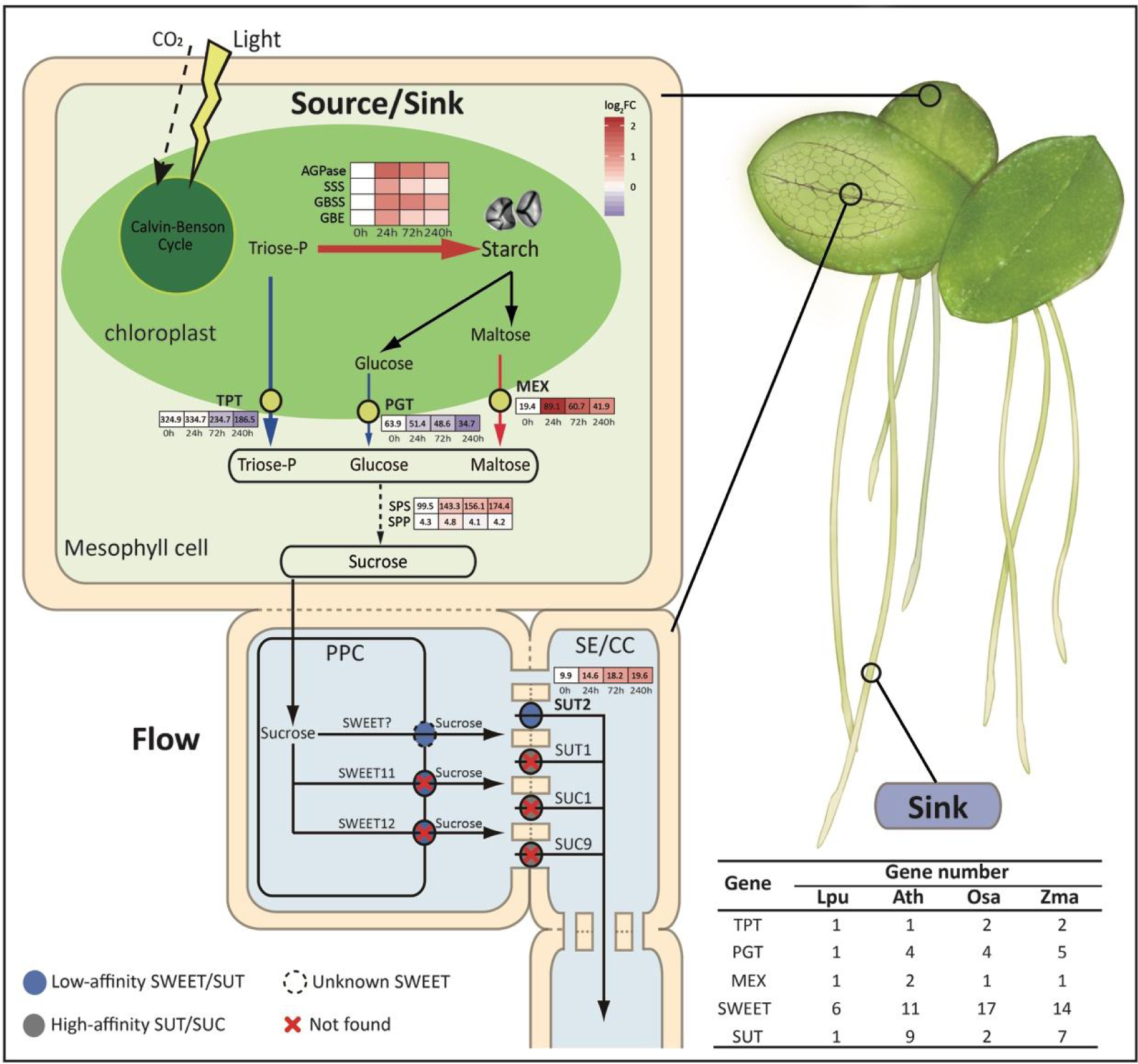
The sugar biosynthesis and transportation in *Landoltia punctate* under LC treatment. The source, flow, and sink in *Landoltia punctata* are represented in light green, light yellow, and light purple, respectively. The heat-maps show the expression profiles of genes involved in starch synthesis, and transportation of triose-P, glucose, and maltose, and sucrose. The numbers in the boxes are the FPKM values. Color of the boxes indicate log_2_FC, where FC was fold changes in expression level comparing with 0 d. The thickness and length of the arrows represents strength of sugar flux. Red, up-regulated expression; blue, down-regulated expression; numbers in the brackets, gene numbers of transporter proteins. SE/CC, the sieve element/companion cell complex.

#### 2.4.3 Weak “flow” in duckweed

The volume of the “flow” in duckweed is affected by the quantity of the sucrose and the efficiency of the transporter. Sucrose content in duckweed is normally < 1.1 mg g^-1^ FW, much lower than that in corn and rice (Fig. S18). Sucrose synthesis is mainly regulated by two enzymes, sucrose phosphate synthase (SPS) and sucrose phosphate phosphatase (SPP). SPS is a reversible rate-limiting enzyme, which catalyzes the synthesis of sucrose-6P using UDP-glucose and fructose-6P and also reversely catalyzes the degradation of sucrose-6P ^52^. The number of *SPS*genes (4) in duckweed is lowercompared with that in rice, corn, and cassava (Data S16a). *SPP*, more importantly, has only one copy and very low expression level (FPKM values <5), resulting in the accumulation of substrate sucrose-6P, then promoting *SPS*to catalyze sucrose-6P degradation reversely. Under LC treatment, the expression level of *SPS*, which FPKM value was already higher than 84 in the beginning (0h), increased significantly (Log_2_FC =1.26 for 240h vs 0h), leading the further enhancement of the degradation of sucrose -6P, and finally resulting in a very low sucrose concentration in the cytoplasm (Data S16b and Fig. S18). Thus, duckweed has an extremely weak ability to synthesize sucrose, and has a low sucrose content in its cell cytoplasm, that is, a low volume of “flow”.

Sucrose transporters (*SUT*s) and hexose and sucrose transporters (*SWEET*s) are responsible for long-distance transport of sucrose to non-photosynthetic organs. Between the two, SUTs are the most important transporter. The duckweed possesses only one *SUT*gene (*LpSUT*) and 6 types of *SWEET*genes (Fig. 4, Data S4 and S6). Compared with Arabidopsis that posesses 9 *SUT*genes, with 4 of which are highly affinitive, the SUT protein in duckweed might be of low affinity. The extended N-terminus of LpSUT lowers its affinity to sucrose, highly similar to SUT2 in Arabidopsis, the sucrose transporter with the lowest affinity ^53^ (Fig. S17). We also observed a very low expression level in *LpSUT*(FPKM values 9.9–19.6) (Data S6). On the other hand, duckweed contains much fewer copies of *SWEET*genes than those of Arabidopsis and rice. Duckweed also lacks the homologs of *SWEET11* and *SWEET12*, the key sucrose efflux transporters ^54^ (Data S4). At the transcriptional level, the expressions of *SWEET*genes in duckweed were also very low (FPKM values <30) (Data S5). Therefore, the number of genes and expression levels of SUT and SWEET showed a weak “flow” ability of the plant. Impressively, the number of genes regulating sucrose transportation are reduced and only the ones with low affinity remained. Due to the reduced gene number and weakened protein activities, sucrose metabolism was markedly suppressed in aspects of synthesis and transportation.

The low sucrose concentration, low sucrose synthesis, and low transport capacity resulted in a weak “flow” in duckweed. LC treatment highly stimulated starch accumulation in the chloroplast and further reduced its sugar flow ability, turning the “source” frond into a “sink” organ (Fig. 4). Therefore, duckweed is an unusual and interesting system in which sources and sinks are spatially organized together, different from the interdependent compartmentation of sinks and sources in other higher plants.

## 3 Conclusions

In the past 50 years, the wide application of green revolution technology resulted in an extraordinary achievement in world staple crop production, especially with the sharp increase in crops harvest index (grain-straw ratio) from 0.3 to 0.5 ^110^. Nowadays, it is very difficult to further increase the harvest index because only certain parts of the crop can be used for harvesting. By contrast, the harvest index of duckweed is nearly 1.0 because of its high starch content and low lignocellulose contents (∼5.8%), especially lignin content (∼1.1%) (Fig. 2). Thus, the whole duckweed can be harvested and used completely. Furthermore, differing from the reproductive growth of cereal crops, duckweed production ability depends on vegetative growth and avoids the time-consuming phase of organ development and differentiation, as well as the fragile stage of sexual reproduction (such as flowering, pollination, etc.) Therefore, the starch productivity of duckweed is considerably higher and more stable than that of staple crops.

The extensive use of green revolution varieties (GRVs) brings excessive consumption and waste of fertilizer and has brought upon serious environmental problems. Since GRV lodging resistance is enhanced by relative insensitivity to nitrogen, GRVs are associated with reduced NUE ^111^. Our results demonstrated that duckweed efficiently assimilated carbon under LC treatment without being supplied with any exogenous nitrogen and phosphorus. The NUE of duckweed reached 144.4 kg biomass kg^-1^ N, much higher than those of maize, rice, and wheat ^112^. In the post green revolution era, an important research and development direction of agriculture is to improve the NUE of crops, starch production using duckweed is a good choice. Meanwhile, LC treatment increased duckweed’s absorption of CO_2_, reducing the emission of greenhouse gas.

The starch content of other duckweed species such as *Spirodelapolyrhiza* and *Lemnaminor* can also reach 45.68-57.23% using this technology, proving its universal applicability for efficient starch production in duckweeds (Fig. S19). Meanwhile, a pilot scale was carried out beside Dianchi Lake, southwest of Kunming (E 102°47′, N 24°51′). The starch content in duckweeds cultivated there can also reach 45.9±3.5% (d.b.) in 4 days, and their starch productivity can reach 36.5 t ha-1 y-1 (Table. S17). Thus, LC treatment has a great potential in practical applications.

To our knowledge, this study is the first report of a simple and environmentally friendly technology for starch production using duckweed. This technology is easy to operate and liable to achieve agricultural industrialization. This work demonstrated that duckweed could be the next generation starch crop and an ideal model plant for starch metabolism research.

## Materials and methods

### 1 Plant material

*Landoltia punctata* strain 0202 was originally obtained at Xinjin, China (N 30°24′46.74″, E 103°48′34.08″) and stored in Chengdu Institute of Biology, Chinese Academy of Sciences (Chengdu, China). The stored duckweed pre-cultured in 1/5 Hoagland medium ^55^ in containers (23×14×4.5 cm^3^) for 7−10 days under a 16 h/8 h (light/dark) photoperiod at 25 °C/15 °C with a light intensity of 110 μmol photons m^-2^ s^-1^ in a greenhouse.

### 2 Cultivation of duckweed with nutrient limitation and/or elevated CO_2_ level

The cultivation of duckweed was conducted in 500 ml beakers (90 mm outer diameter ×120 mm height) containing 500 mL medium (1/5 Hoagland medium or deionized water) with an initial inoculation of 1 g fresh pre-cultivated duckweed. This experiment incorporates three treatment conditions: nutrient limitation (L), cultivating duckweed in deionized water; elevated CO2 level (C), supplying CO_2_ to 2500±100 ppm (Table S1) ^56,57^; and the combination of L and C (LC). With the three variables listed above, all duckweed was cultivated for 10 days under a 24 h/0 h (light/dark) photoperiod at 25 °C with a light intensity of 110 μmol photons m^-2^ s^-1^. Fresh duckweed of 0.5g from each sample were collected at 0 h, 2 h, 5 h, 9 h, 24 h, 48 h, 72 h, 120 h, 168 h, and 240 h and snap-frozen immediately in liquid nitrogen. Samples were stored at −80 °C for subsequent biophysiological and biochemical analysis, transcriptome sequencing, and/or whole genomic bisulfite sequencing (WGBS). Three biological replicates were performed to acquire the mean values for all data in the experiment, except for WGBS.

On a pilot scale, duckweed was treated under LC condition for one month (from 14 February to 13 March, 2014) beside Dianchi Lake, southwest of Kunming, E 102°47′, N 24°51′). Approximately 4.0 kg fresh duckweed was transferred into 3.1×4.5×0.4 m^3^ (W×L×D) tanks filled with tap water in a greenhouse where CO_2_ was aerated to a concentration of 2500±100 ppm. Duckweed was cultivated at 20−30 °C with sunlight in the daytime and fluorescent lamp at night for 4 days (Table S17).

### 3 Light microscopy and transmission electron microscopy (TEM)

Duckweed fronds in the treatment and control went through processes of fixation, embedding, and dehydration as described previously ^58^. Semithin sections were stained with 0.2% (w/v) KI/I_2_ solution and observed under a Motic BA210 microscope equipped with a digital camera (Fig. S1). Ultrathin sections (80 nm thick) of duckweed were cut with an ultramicrotome (Leica EM UC7, Leica) and observed under a transmission electron microscope (Hitachi H-7650TEM, Japan) ^59^. Images were processed (sharpened, brightened, and contrast adjusted) and assembled using Photoshop CS6 (Adobe).

### 4 Gene family analysis

The 8 species we collected protein sets from to conduct gene family analysis are listed as follow: Klebsormidiumflaccidum^60^,Zosteramarina^61^,Arabidopsisthaliana ^62^, Oryzasativa japonica ^63^, Zeamays^64^, Spirodelapolyrhiza^65^, Landoltiapunctata, and Lemna minor ^66^. Using the results from ‘all-versus-all’ BLASTP (e-value threshold 1 × 10^-3^) comparison, OrthoMCL ^67^ (mcl –I 1.5) was used to delineate gene families.

### 5 Transcriptome analysis

Total RNA was extracted using OMEGA^TM^ Plant DNA/RNA Kit (OMEGA, USA) following the manual. Genomic DNA was removed by DNase I (Fermentas, USA). RNA concentrations, quality, and integrity number (RIN) were measured with Agilent 2100 Bioanalyzer (Agilent, USA). Pair-end sequencing (2×150 bp) using Illumina HiSeq 2500 platform at Mega Genomics Co (Beijing, China) was conducted on libraries.

All raw sequences were evaluated by FastQC_v0.11.3 (http://www.bioinformatics.babraham.ac.uk/projects/fastqc/) where low quality sequences (reads with adapter, ambiguous base ‘N’, or low-quality scores) were filtered out. The high quality clean reads were aligned to ribosomal RNA database (rRNA) using Bowtie2 ^68^ to remove rRNA reads. Then, Hisat2-2.0.5 ^49^ was used to align the reads to the reference genome, and Stringtie-1.3 ^69^ was used to calculate FPKM values. Significant differences in expression levels were evaluated using Ballgown_2.6.0 ^70^ (p value ≤ 0.05, |log_2_(fold change)| ≥ 0.58).

Twenty differentially expressed genes (DEGs) were validated by quantitative reverse transcription-PCR (qRT-PCR) analysis. Total RNA was extracted from backup samples for transcriptome analysis using Eastep^®^ Super Total RNA Extraction Kit (Promega, USA). Reverse transcription was performed using GoScript^TM^ Reverse Transcription System (Promega, USA). qRT-PCR was performed using UltraSYBR Mixture (Cwbiotech, China) with CFX Connect Real-Time PCR System (Bio-Rad). *Actin* was used as the reference gene. Primers were listed in Table S13. Three technical replicates of qRT-PCR reaction was conducted for each sample.

### 6 Quantification of key genes expression

Expression of key genes involved in CO_2_ fixation, carbon concentration, and starch synthesis (*PEPC*, *Rubisco*, *UGPase*, *AGPase*, *SSS*, and *GBSS*) was quantified by qRT-PCR. qRT-PCR followed the protocol as described above with *Actin*as the reference gene where three technical replicates were again performed. Primers were listed in Table S14.

### 7 Subcellular localizations of enzymes and translocators

To explore the subcellular localization of target proteins, the coding regions of corresponding genes in *Landoltia punctata* were first cloned independently into the binary vector pCAMBIA2300-GFP, which were then transformed independently into the *Agrobacterium* strain GV3101. The suspension cells of *Lemna gibba* were infiltrated with *Agrobacterium* strain GV3101 that carried either GFP-fused C-terminus or N-terminus of the target protein, or empty vector (control) ^71^. After infiltrating the suspension cells with *Agrobacterium* for 3 days, protoplasts were generated by Cellulase R10 and Macerozyme R10 (Yakult Pharmaceutical Ind. Co., Ltd., Japan) digestion ^72^. All of the fluorescence signals were detected using a confocal laser scanning microscope (Leica TCS SP8). The excitation/emission spectra were 488/493 to 598 for GFP and 633/647 to 721 for chlorophyll auto-fluorescence.

### 8 DNA methylation analysis

Following the manufacturer’s instructions of Acegen Bisulfite-Seq Library Prep Kit (Acegen, Shenzhen, China Cat. #BS0311-48), a WGBS library was constructed using 500 ng of purified genomic DNA spiked with 0.1% (w/w) unmethylated Lambda DNA (Promega, Madison, WI). Briefly, DNA was sonicated (Covaris) to a mean fragment size distribution of 200–400 bp. Fragmented DNA was end-repaired, 5’-phosphorylated, 3’-dA-tailed, and ligated to adapters. The adapter-ligated DNA molecules were purified using 1× Agencourt AMPure XP magnetic beads and were subjected to bisulfite conversion using ZYMO EZ DNA Methylation-Gold Kit (Zymo, Cat. #D5005). Libraries were then amplified by PCR using 20 μL of bisulfite-converted DNA molecules, 25 μL of KAPA HiFi HotStart Uracil+ ReadyMix, and 5 μL of 8-bp index primers each with a final concentration of 1 μM. PCR was performed under cycle conditions of an initial denaturation at 98 °C for 1 min, 10 cycles of 98 °C for 15 s, 60 °C for 30 s, and 72 °C for 30 s, and extension for 1 min at 72 °C. Constructed WGBS libraries were then analyzed by Agilent 2100 Bioanalyzer and quantified by a Qubit fluorometer with Quant-iT dsDNA HS Assay Kit (Invitrogen).

Pair-end sequencing (2 × 150 bp) was done on WGBS libraries using the Illumina HiSeq X Ten platform at Mega Genomics Co. (Beijing, China). All raw sequences were evaluated by FastQC_v0.11.3 (http://www.bioinformatics.babraham.ac.uk/projects/fastqc/) where low-quality sequences (reads with adapter, 5% ambiguous base ‘N’, or low quality scores) were filtered out. Clean reads were aligned against the *Landoltia punctata* genome using the BSMAP 2.90 ^73^ under default parameters. Identification of methylated cytosine positions for each sample was performed independently in accordance with previous study ^74^. The CG, CHG, and CHH methylation rates of genes were counted using AWK script.

The samples under methylation inhibitor treatment were sent to Basebio Co. (Chengdu, China) for WGBS. The WGBS library construction, quality control, and sequencing (Illumina HiSeq 4000) were executed with accordance to methods described above. Clean reads were aligned against the reference genome using WALT ^75^ with default parameters and then de-duplicated before downstream analysis. MethPipe ^76^ was used to identify sites of methylation, where at least five reads containing cytosine considered. The binomial test was performed for each cytosine base to check the methylated cytosine site (mC) with false discovery rate ≤ 0.05. The methylation level (ML) of each target region was calculated with Eq. (1) using ViewBS ^77^ as follows:

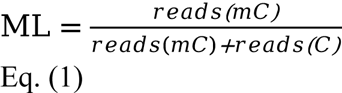

### 9 Analytical methods

#### 9.1 Composition analysis

Prior to analysis, duckweed was dried to constant weight at 60 °C and milled. Structural carbohydrates, including glucan, xylan, galactan, arabinan, mannan, lignin, and ash were determined according to the method recommended by National Renewable Energy Laboratory, USA ^78^. Starch content was determined according to the procedure based on hydrolysis of duckweed using HCl, as described previously ^17^. Cellulose content was calculated by subtracting starch from glucan. Xylan, galactan, arabinan, and mannan together was considered hemicellulose. Lipid was extracted using diethyl ether with a Soxtec system with reference to AOAC 920.39 B (http://down.foodmate.net/standard/sort/10/25070.html). Total Kjeldahl nitrogen (TKN) was measured by a FOSS KJ2200 System (FOSS Corp., Denmark). Protein content was calculated by TKN content times conventional factor (6.25). Pectin was extracted according to the method described by ^79^ and ^80^, and determined according to the method described by ^81^ using GalUA (Sigma-Aldrich) as a standard. Elements of carbon and nitrogen were determined by an elemental analyzer (Vario EL Cube; Elementar Analysensysteme GmbH, Germany).

#### 9.2 Enzyme activity assay

Fresh duckweed (0.5 g) was homogenized in 5 ml precooled enzyme extracting solution (100 mM Tricine-NaOH (pH 8.0), 8 mM MgCl_2_, 2 mM EDTA, 50 mM 2-mercaptoethanol, 12.5% (v/v) glycerol, and 5% (w/v) insoluble polyvinylpyrrolidone-40) ^82^. The homogenate was centrifuged at 13,400 *g*for 10 min at 4 °C. Then, the supernatant was used to measure the enzyme activities. Activities of AGPase (EC 2.7.7.27) and starch synthase (SSS, EC 2.4.1.21; GBSS, EC 2.4.1.242) were analyzed using the methods described by Nakamura, et al. ^82^. The activities of the two enzymes were tested by the change of NADH at 340 nm measured by a microplate reader (Thermo Scientific Varioskan Flash, Thermo Fisher Scientific Inc., USA). Activities of α-amylase (EC 3.2.1.1) and β-amylase (EC 3.2.1.2) were estimated following the method ^17^. Rubisco (EC 4.1.1.39) activity was tested as described by Sharkey, et al. ^83^ using a spectrophotometric diagnostic kit (Suzhou Comin Biotechnology Co., Ltd., Suzhou, China). PEPC (EC 4.1.1.31), NADP-MDH (EC 1.1.1.82), and malic enzyme (ME, EC 1.1.1.39) activities were assayed according to Gonzalez, et al. ^84^ and Johnson and Hatch ^85^ using a spectrophotometric diagnostic kit (Suzhou Comin Biotechnology Co., Ltd., Suzhou, China).

#### 9.3 Determination of intracellular sucrose

Sucrose was extracted according to the method described by ^86^. Duckweed was dried to constant weight at 60 °C and then powdered using a pulverizer. Then, 50 mg duckweed powder was suspended in 1 ml deionized water and sonicated for 30 min. The mixture was incubated at 80 °C for 1 h with intermittent shaking every 5 min. The sample was centrifuged at 13,400 *g*for 20 min at 4 °C. The supernatant was store at −20 °C. The residue was re-suspended in 1 ml deionized water to a second-round extraction. The supernatants from two rounds of extraction were pooled and filtered through a 0.45-μm-pore size filter.

Sucrose content was determined using a high-performance liquid chromatography (HPLC) system (Thermo 2795, Thermo Corp.) equipped with an evaporative light scattering detector (ELSD) (All-Tech ELSD6000, All-tech., Corp.). Samples were separated by Aminex HPX-87P column (300 × 7.8 mm) at 79 °C using ultrapure water as mobile phase at 0.6 ml min^-1^. Analytical pure sucrose (AR) was applied as a standard.

#### 9.4 Determination of Glycerate 3-P (3-PGA)

3-PGA content was measured by an enzymatic assay described by ^87^. First, 3-PGA was abstracted using precooled methanol/chloroform (1:1, v/v). Then, the endogenous enzymes in the samples were heat-inactivated at 70 °C for 10 min. Next, 20 μL resulting extract was added into 980 μL reagent (0.1 M Tris-HCl (pH 7.6), 5mM MgCl_2_, 40 μM NADH, 2 mM ATP, 6 units PGK and 3 units GAPDH). The reaction was kept at 25 °C for 20 min. Finally, the samples were tested at 340 nm using spectrophotometer. In control, 20 μL methanol/chloroform (1:1, v/v) instead of extract was added into reagent.

### 10 Identification of transcription factors in starch biosynthetic pathway in *Landoltia punctata*by gene co-expression analysis

The transcriptome data of nutrient limitation and elevated CO_2_ level (2500±100 ppm) (LC0, LC1, LC3) was used for co-expression analysis ^88^. The transcription factor library of *Landoltia punctata* were constructed with iTAK ^89^. The absolute value of the Pearson correlation coefficient between the key genes of starch biosynthetic pathway (*AGPase,SSS,GBSS,GBE*) and TFs was calculated using Hmisc package ^90^. The top 30 of absolute value of the Pearson correlation coefficient greater than 0.8 were listed in Data S17 with FPKM value of TFs.

## Acknowledgments

We thank Zhongyan Wang for technical support. We thank Wan Xiong for language editing. We also thank Ping Mao for providing information. This research was supported by Innovation Academy for Seed Design, CAS; National Aquatic Biological Resource Center (NABRC); the National Natural Science for General Foundation of China (31770395); Key deployment projects of Chinese Academy of Sciences (ZDRW-ZS-2017-2-1); and Biological Resources Programme, Chinese Academy of Sciences (KFJ-BRP-008).

## Author contributions

H.Z, Y.F, and S.W conceived and designed the study. H.Z, Y.F, and S.C supervised and funded the study. Y.F, L.G, Y.X, Y.J, A.D, X.T, Z.L, S.C, L.T, Y.Z, J.L, F.Z, and Y.H performed the experiments. Y.F, L.G, Y.X, and Y.Z performed the field work. S.W, Y.X, Y.D, A.D, X.T, Y.C, Z.Y, and L.C performed bioinformatics analysis. H.Z, Y.F, L.G, S.W, Y.X, Y.J, A.D, X.T, Y.D, Z.L, K.H, L.T, Z.Y, L.C, P.Z, Z.G, F.Z, and Q.Z contributed to the discussion and important intellectual content. Y.F, H.Z, L.G, S.W, L.Z, A.D, X.T, Z.L, K.H, S.C, P.Z, Z.G, and Q.Z wrote and revised the manuscript.

## Data and materials availability

All data are available in the manuscript, the supplements, or at publicly accessible repositories. Raw reads of whole genome sequencing of *Landoltia punctata* 0202 have been deposited at NCBI under BioProject ID: PRJNA546087. The whole genome data of other species used in this study are available in the supplementary materials. All transcriptomes have been uploaded to NCBI under BioProject ID: PRJNA672224. All epigenetic data have been deposited in NCBI under BioProject ID: PRJNA673253. The duckweeds are available from the Duckweed Resource Bank in Chengdu Institute of Biology, Chinese Academy of Sciences.

