## Supplementary Materials for "Artificially cultivated duckweed: a high-efficiency starch producer"

### Supporting Information

#### Supplementary Text

1 Transcriptomes

1.1 RNA Sequencing

7.2 Gb raw bases were acquired for each duckweed sample (Table S2), with approximately 7 Gb of clean bases. The Q20s of bases were above 97.2%, indicating that the sequences could be used for further analysis.

1.2 Validation of RNA Sequencing

Twenty DEGs were selected to validate RNA sequencing by qRT-PCR. Among the selected DEGs, the expression patterns of 19 genes in qRT-PCR were similar to those in RNA sequencing (Fig. S4), suggesting an overall consistency between RNA sequencing data and qRT-PCR results. Therefore, the RNA-sequencing in this study was reliable.

2 Quantification of key genes expression

Expression of key genes involved in $CO_2$ fixation, carbon concentration, and starch synthesis (*PEPC*, *Rubisco*, *UGPase*, *AGPase*, *SSS*, and *GBSS*) was quantified by qRT-PCR. Results from qRT-PCR of key genes (Fig. S5) were similar to those of FPKM values from RNA sequencing, showing an increase of the expression level of most key genes induced by LC treatment.

3 Subcellular localizations of enzymes and translocators

We investigated the subcellular locations of AGPase (10015507), GBSS (10021837), SSS (10010699), UGPase (10020333), plastidic glucose translocator (PGT), maltose transporter (MEX), triose phosphate/phosphate translocator (TPT), sucrose transporters (SUT), hexose and sucrose transporter SWEET1 (10012584), and SWEET7B (10002893) in *Landoltia punctata*. To this end, GFP were fused to the N-terminus of SUT protein and to the C-terminus of the rest of the proteins. These proteins were then assessed for their subcellular localization in duckweed protoplast cells.

Transient expression of AGPase-, GBSS-, and SSS-GFP fusion protein in protoplasts revealed their co-localization with chlorophyll, which confirms their presence in chloroplasts and their participation in starch biosynthesis (Fig. S6 A-C). Transient expressions of PGT-, MEX-, TPT-GFP fusion proteins also confirmed the transporters localized mainly on chloroplasts. Thus, the three transporters in chloroplasts might work for sugar transport (Fig. S6 D-F) (*1*). Specially, transient expression of UGPase-GFP fusion protein showed it localized mainly in cytoplasm, suggesting UGPase as a cytosolic type in *Landoltia punctata* (Fig. S6 G).

Transient expression of SWEET1- and SWEET7B-GFP fusion protein in protoplasts revealed that they localized mainly on cell membrane, confirming SWEET transporters' ability to transport sugar on cell membranes (Fig. S7 A-B). The transient expression of sucrose transporter (SUT) in protoplasts showed that GFP-SUT fluorescence is mainly retained in intracellular structures, similar to the SISUT2 subcellular localization in tomato (Fig. S7 C) (*2*). This suggests it might lose its function as a sucrose transporter in *Landoltia punctata*.

<u>4 DNA methylation analysis</u>

WGBS libraries were subjected to pair-end sequencing using Illumina HiSeq X Ten platform. Filtered high quality reads were aligned against the genome of *Landoltia punctata* (Table S5). Identification of methylated cytosine positions for each sample was performed independently.

The global methylation level of *Landoltia punctata* ranged from 11.2% to 13.0%. Primarily, methylation level involving CG sequences was more than 70% (Table S6 and Fig. S20). The methylation levels of different genome regions (exons, introns, and 2 kb-upstream and -downstream of genes) decreased under LC treatment (Fig. S21). In particular, methylation levels of CG sequences in exons, introns, and the 2 kb-upstream of genes decreased dramatically and remained relatively low during the 10-day cultivation under LC treatment (Fig. S22).

<u>5 Universal applicability of LC treatment</u>

Two of other duckweed species, *Spirodela polyrhiza* and *Lemna minor*, also accumulated starch content to over 45% under LC treatment (Fig. S19). In the pilot-scale test when duckweed was harvested every 4 days, starch content and yield reached 45.9 ± 3.5 % and 10.0 ± 1.4 g $m^{-2}$ $d^{-1}$, equivalent to annual starch yield of approximately 36.5 t $ha^{-1}$ (Table S7). This indicated the universal applicability of LC treatment for efficient starch production in duckweeds.

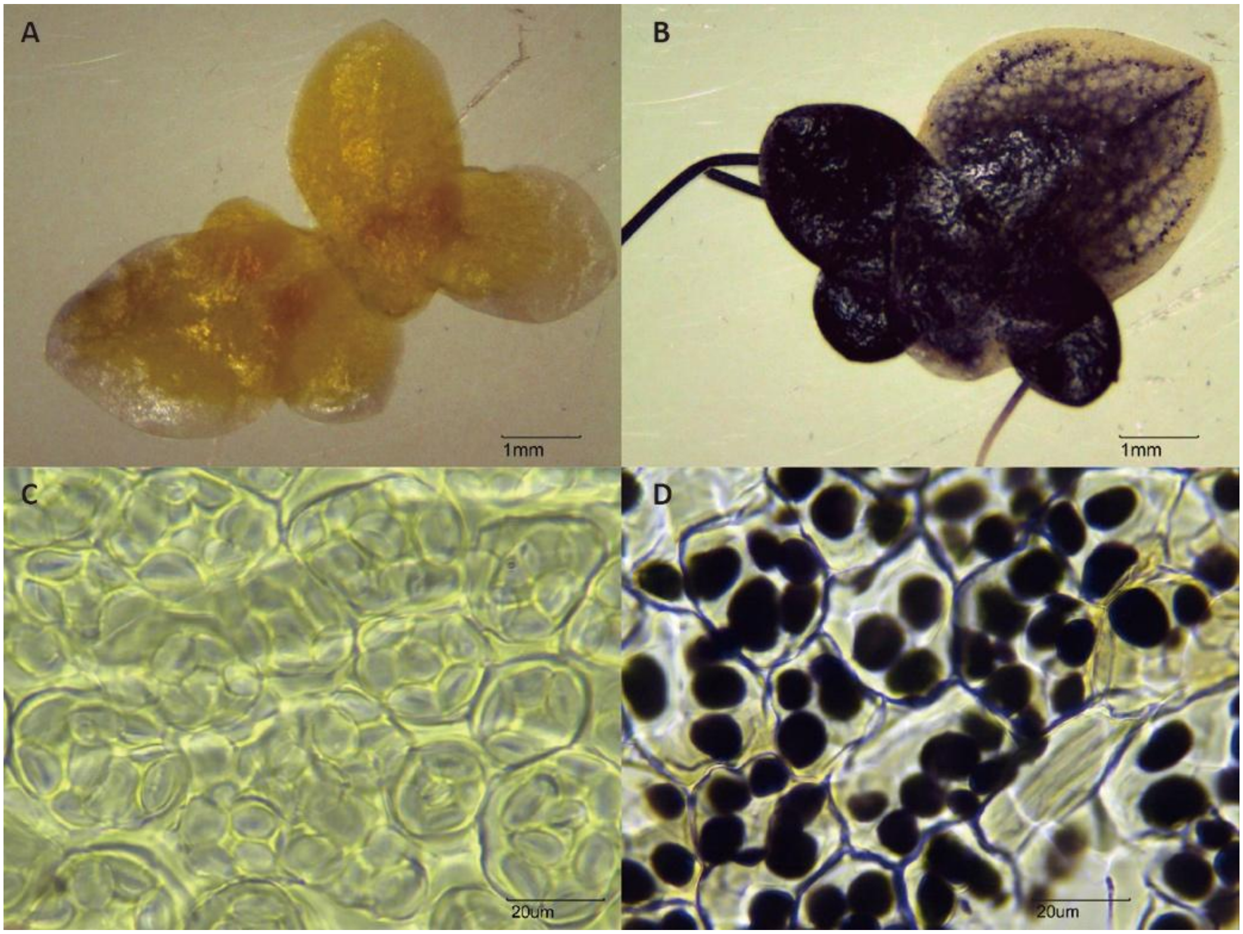


**Fig. S1.**

Iodine staining of starch granules in *Landoltia punctata*.

**A and C**, iodine staining of the whole frond (A) and semithin sections (C) in the control treatment. The fronds were cultivated in 1/5 Hoagland media for 72 h.

**B and D**, iodine staining of the whole frond (B) and semithin sections (D) in the LC treatment. The fronds were cultivated under conditions of nutrient limitation and elevated $CO_2$ level (2500±100 ppm) for 72 h.

Bars, 1 mm in A and B; 20 μm in C and D.

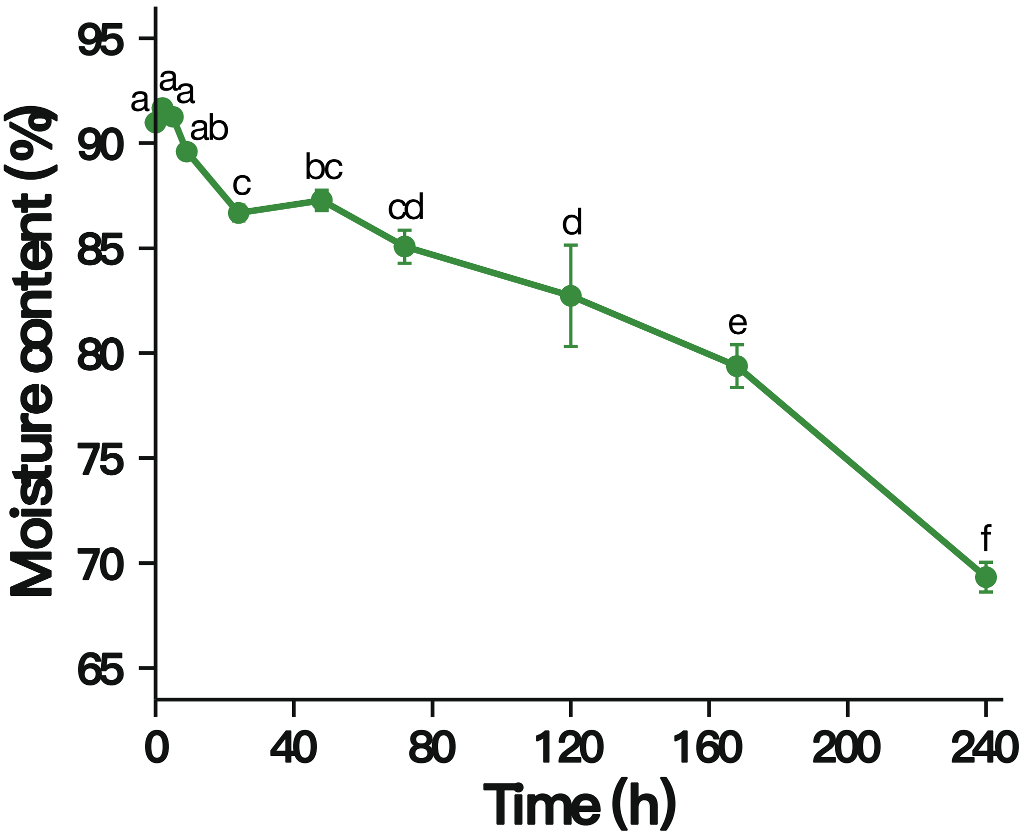


**Fig. S2.**

Changes in moisture content of *Landoltia punctata* under LC treatment.

LC, cultivated under conditions of nutrient limitation and elevated $CO_2$ level (2500±100 ppm).

Letters indicate significant differences among time points, tested by one way ANOVA following Tukey-Kramer test ($p<0.05$).

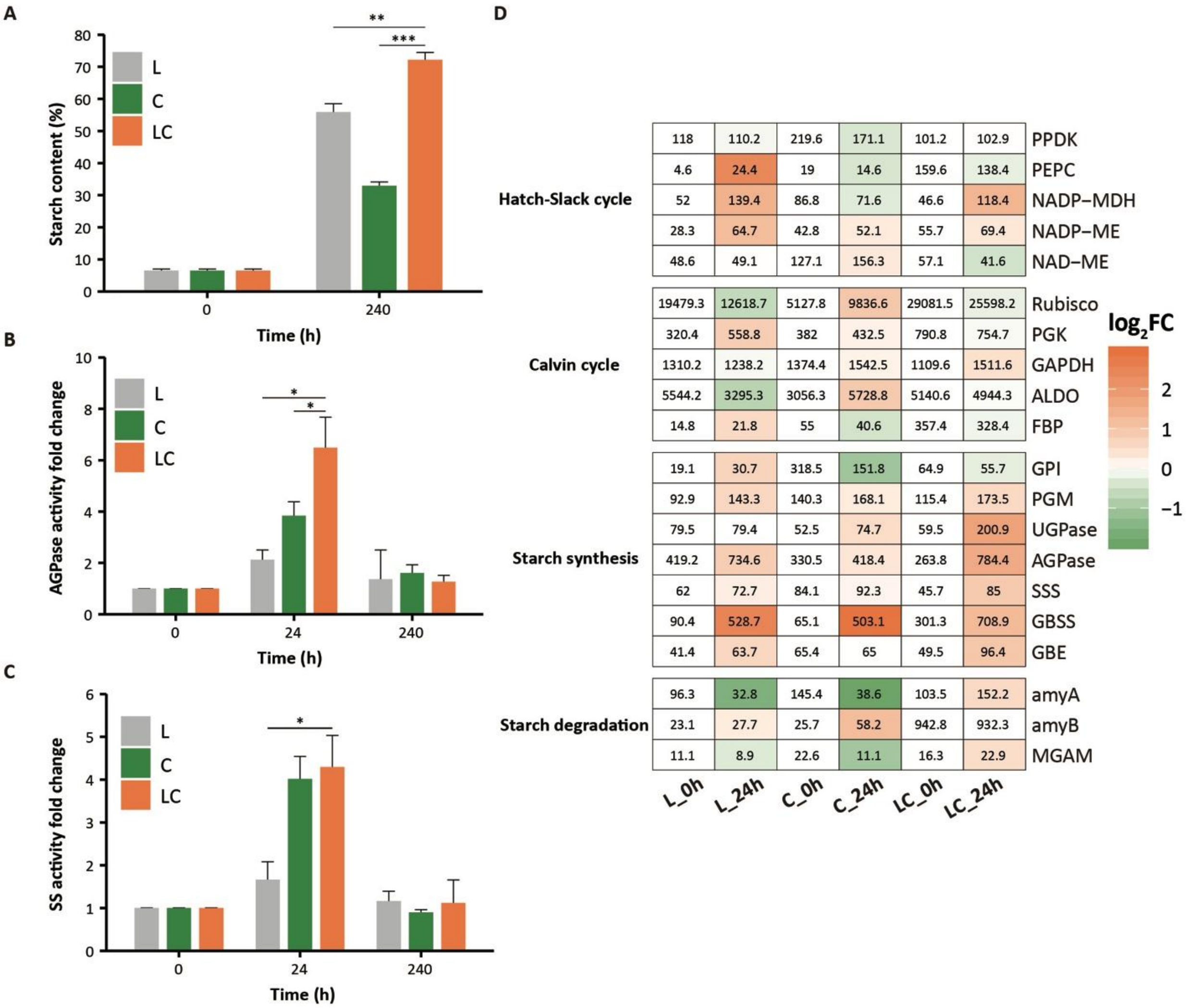


**Fig. S3.**

Starch accumulation and biosynthesis in *Landoltia punctata* under different conditions.

**A**, Starch content.

**B**, Fold change of AGPase activity compared with 0 h.

**C**, Fold change of SS activity compared with 0 h.

**D**, Expression of genes involved in starch biosynthesis under various conditions. Numbers indicate the FPKM values. Colors indicate $log_2FC$ comparisons of expression values at 24 h and 0 h.

L, cultivated under condition of nutrition limitation;

C, cultivated under condition of elevated $CO_2$ level (2500±100 ppm);

LC, cultivated under conditions of nutrient limitation and elevated $CO_2$ level (2500±100 ppm). Error bars are standard deviations measured from three independent cultures. Asterisk indicates statistically significant difference between treatment group and control in the same assay conditions (Student's *t*-test). *, $P<0.05$; **, $P<0.01$; ***, $P<0.001$.

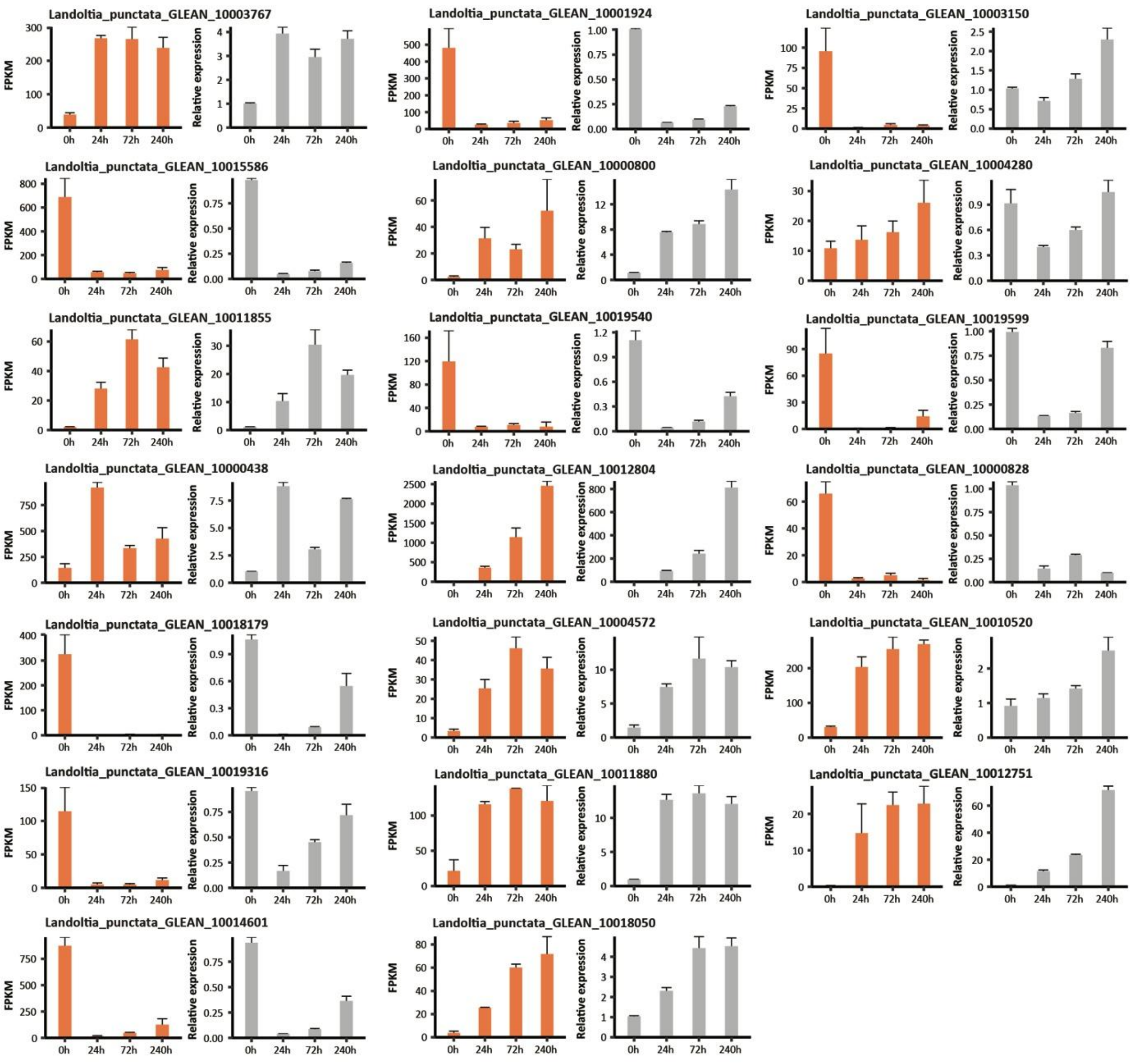


**Fig. S4.**

Validation of RNA-Sequencing by qRT-PCR

For each pair of graphs, the left exhibits results from RNA sequencing while the right exhibits qRT-PCR data. Y-axis of RNA sequencing graphs represent FPKM value, while that of qRT-PCR graphs represent relative expression level, the expression level of the target gene normalized to that of internal control *Actin*.

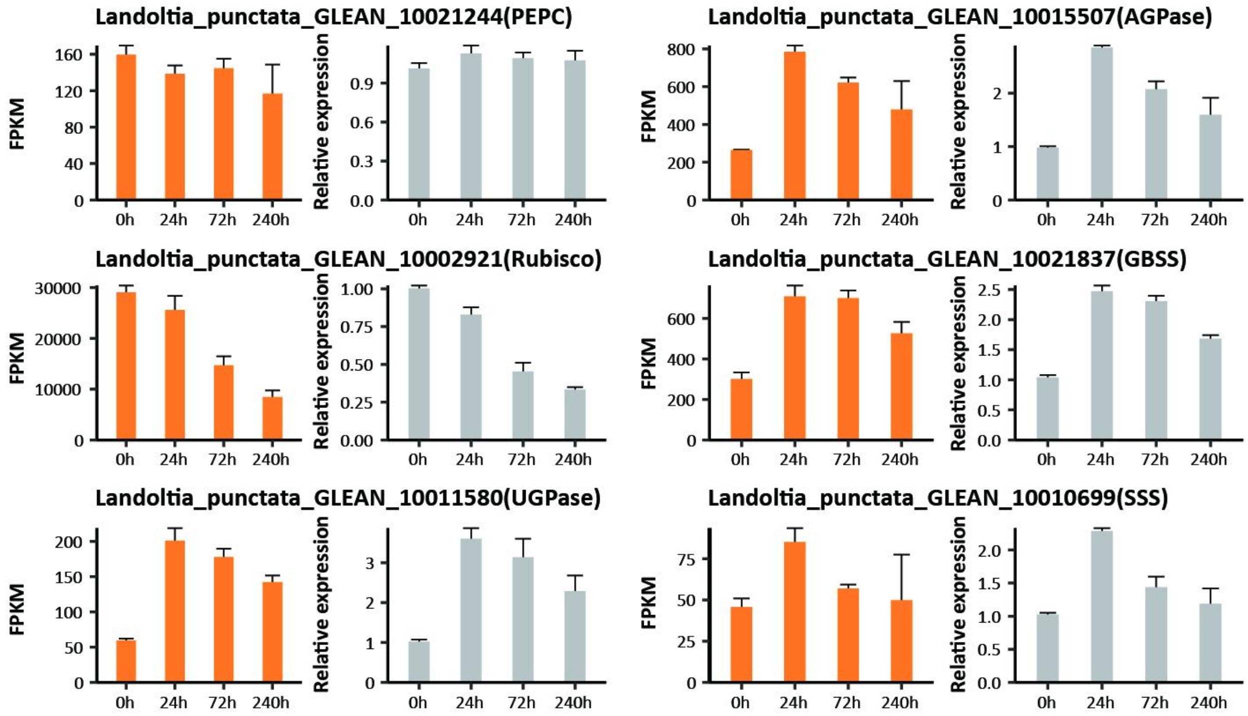


**Fig. S5.**

Expression levels of key genes involved in $CO_2$ fixation, carbon concentration, and starch synthesis as analyzed by qRT-PCR.

For each pair of graphs, the left exhibits result from RNA sequencing while the right exhibits qRT-PCR data. Y-axis of RNA sequencing graphs represent FPKM value, while that of qRT-PCR graphs represent relative expression level, the expression level of the target gene normalized to that of internal control *Actin*.

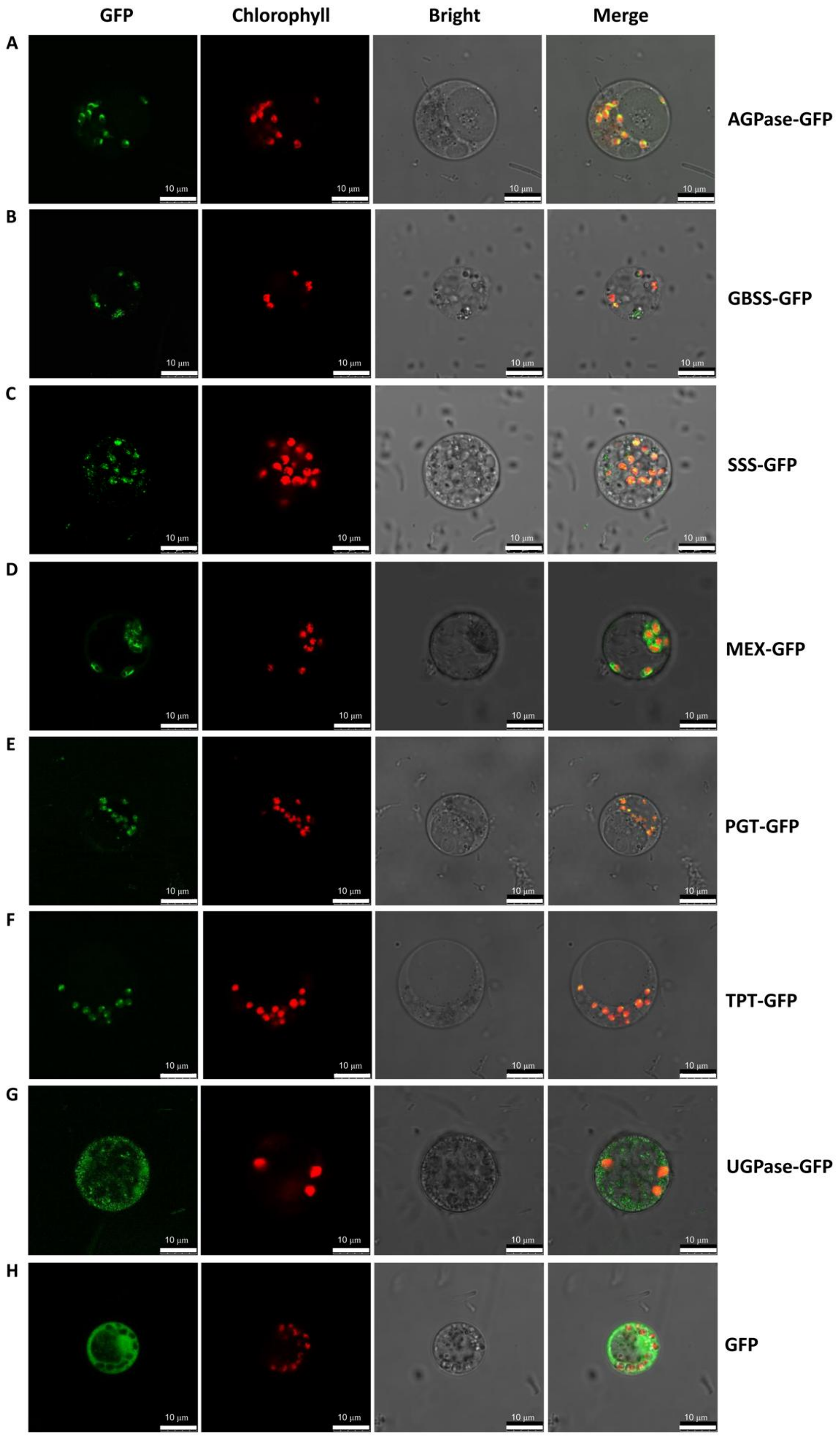

GFP
Chlorophyll
Bright
Merge
A
AGPase-GFP
B
GBSS-GFP
C
SSS-GFP
D
MEX-GFP
E
PGT-GFP
F
TPT-GFP
G
UGPase-GFP
H
GFP
10 μm

**Fig. S6.**

Confocal laser scanning microscopy images of duckweed (*Lemna minor*) protoplast cells showing

Subcellular localization of AGPase (10015507), GBSS (10021837), SSS (10010699), MEX, PG T, TPT, and UGPase (10020333) proteins.

**A-G**, target proteins with GPF-fused C-terminus

**H**, GFP protein, used as control

The green fluorescence signal indicated the target proteins or GFP protein.

The red fluorescence signal indicated the autofluorescence of chloroplasts.

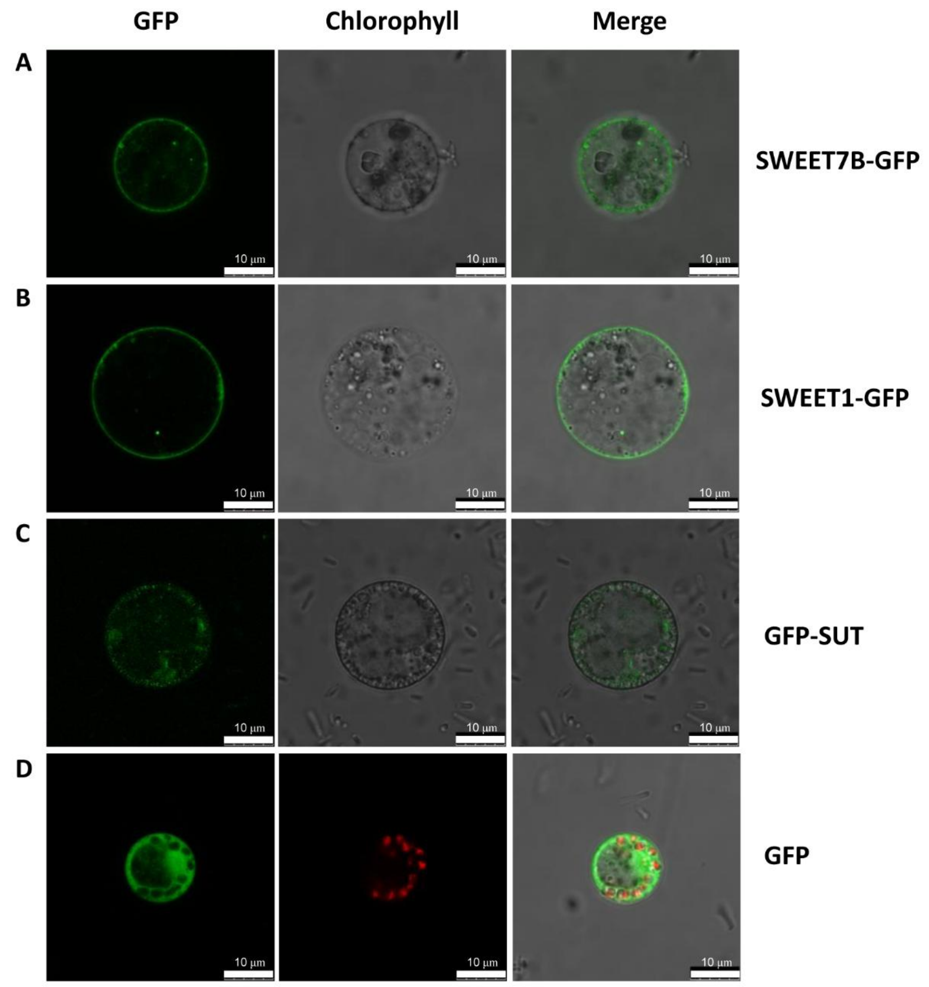


**Fig. S7.**

Confocal laser scanning microscopy images of duckweed (*Lemna minor*) protoplast cells showing

subcellular localization of SWEET (SWEET7B, 10002893; SWEET1, 10012584) and SUT prote ins.

**A and B**, SWEET proteins with GPF-fused C-terminus

**C**, SUT protein with GFP-fused N-terminus

**D**, GFP protein, used as control

The green fluorescence signal indicated the fusion proteins. The red fluorescence signal indicated the autofluorescence of chloroplasts.

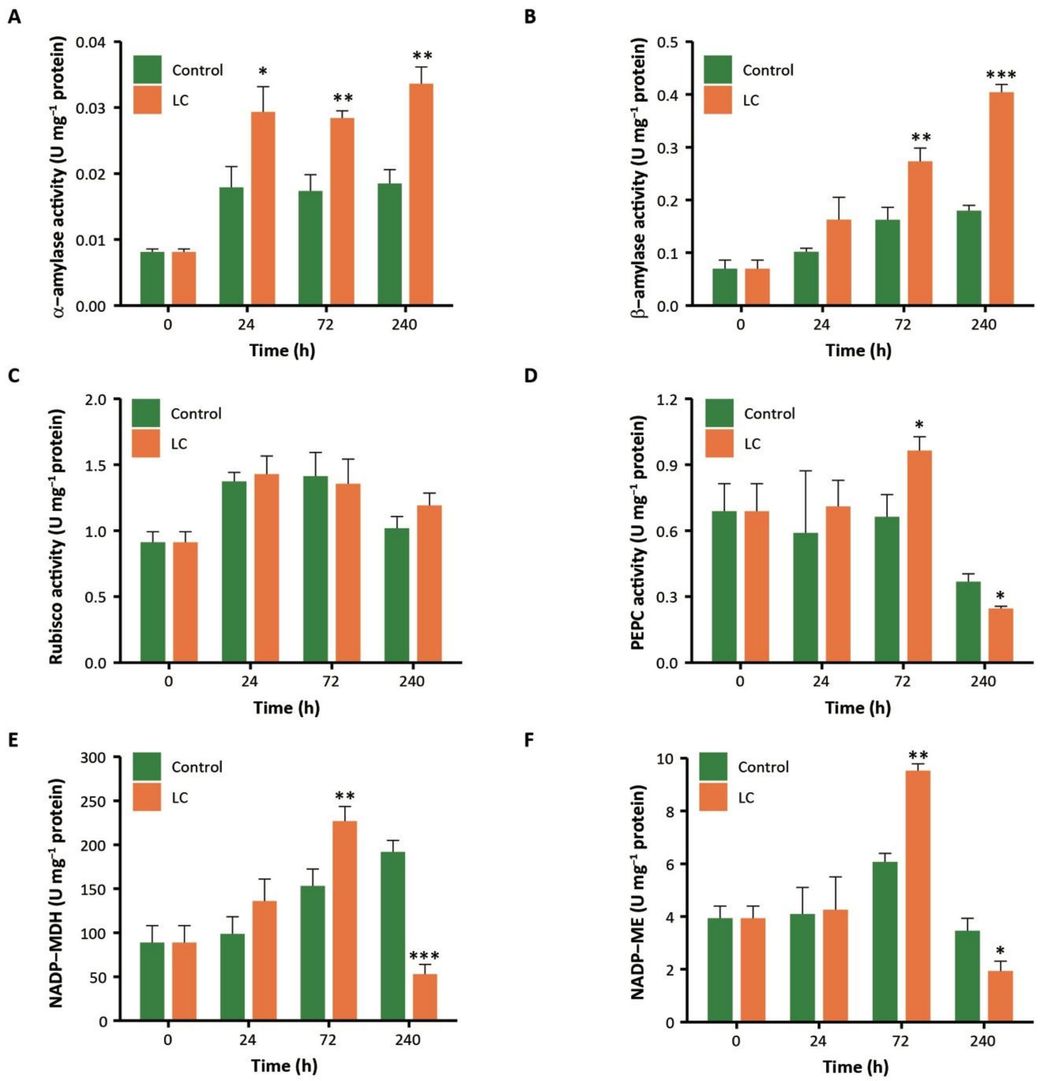


**Fig. S8.**

Activities of (**A**) α-amylase, (**B**) β-amylase, (**C**) Rubisco, (**D**) PEPC, (**E**) NADP-MDH, and (**F**) NADP-ME in *Landoltia punctata*.

LC, cultivated under conditions of nutrient limitation and elevated $CO_2$ level (2500±100 ppm). Control, cultivated in 1/5 Hoagland medium. Rubisco, ribulose-bisphosphate carboxylase; PEPC, phosphoenolpyruvate carboxylase; NADP-MDH, malate dehydrogenase ($NADP^+$); NADP-ME, malate dehydrogenase ($NADP^+$). Error bars show standard deviations measured from three independent cultures. Asterisk indicates statistically significant difference between treatment group and control in the same assay conditions (Student's *t*-test). *, P<0.05; **, P<0.01; ***, P<0.001.

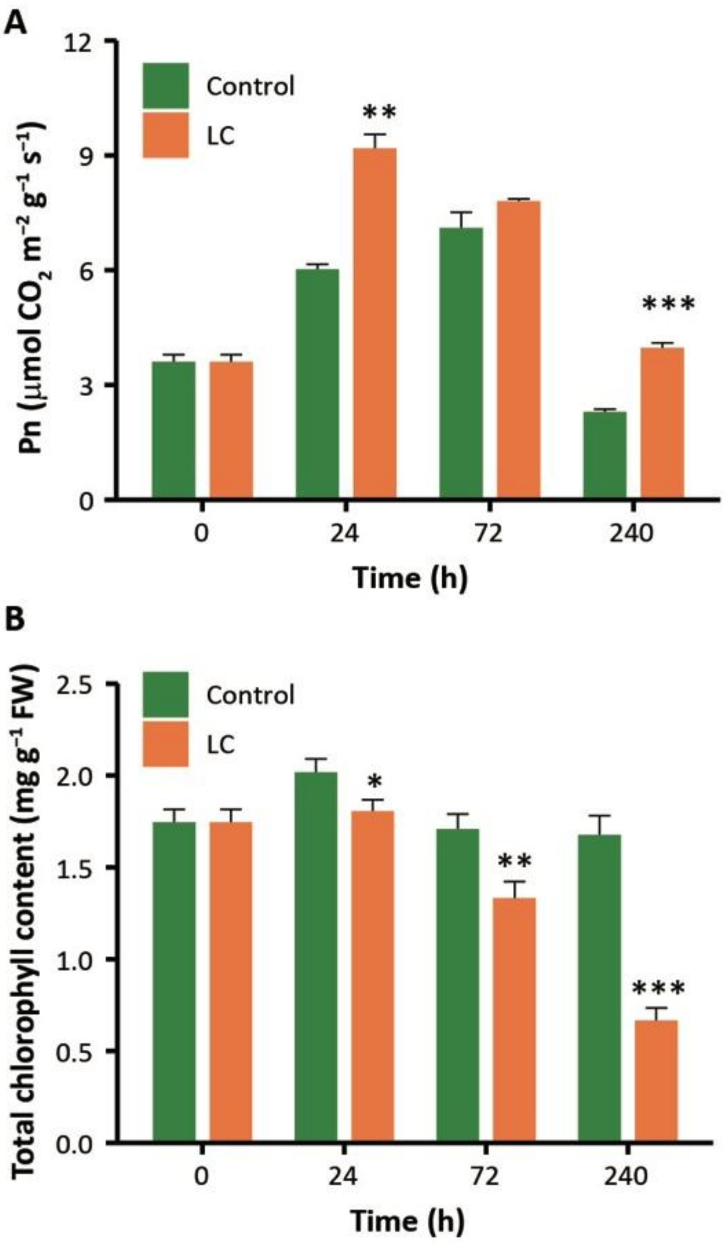


**Fig. S9.**

Net photosynthetic rate (**A**) and total chlorophyll content (**B**) of *Landoltia punctata*.

LC*,* cultivated under conditions of nutrient limitation and elevated $CO_2$ level (2500±100 ppm). Control, cultivated in 1/5 Hoagland medium. Pn, net photosynthetic rate. Error bars represent the SDs measured from three independent cultures. Asterisk indicates statistically significant difference between treatment group and control in the same assay conditions (Student's *t*-test). *, $P<0.05$; **, $P<0.01$; ***, $P<0.001$.

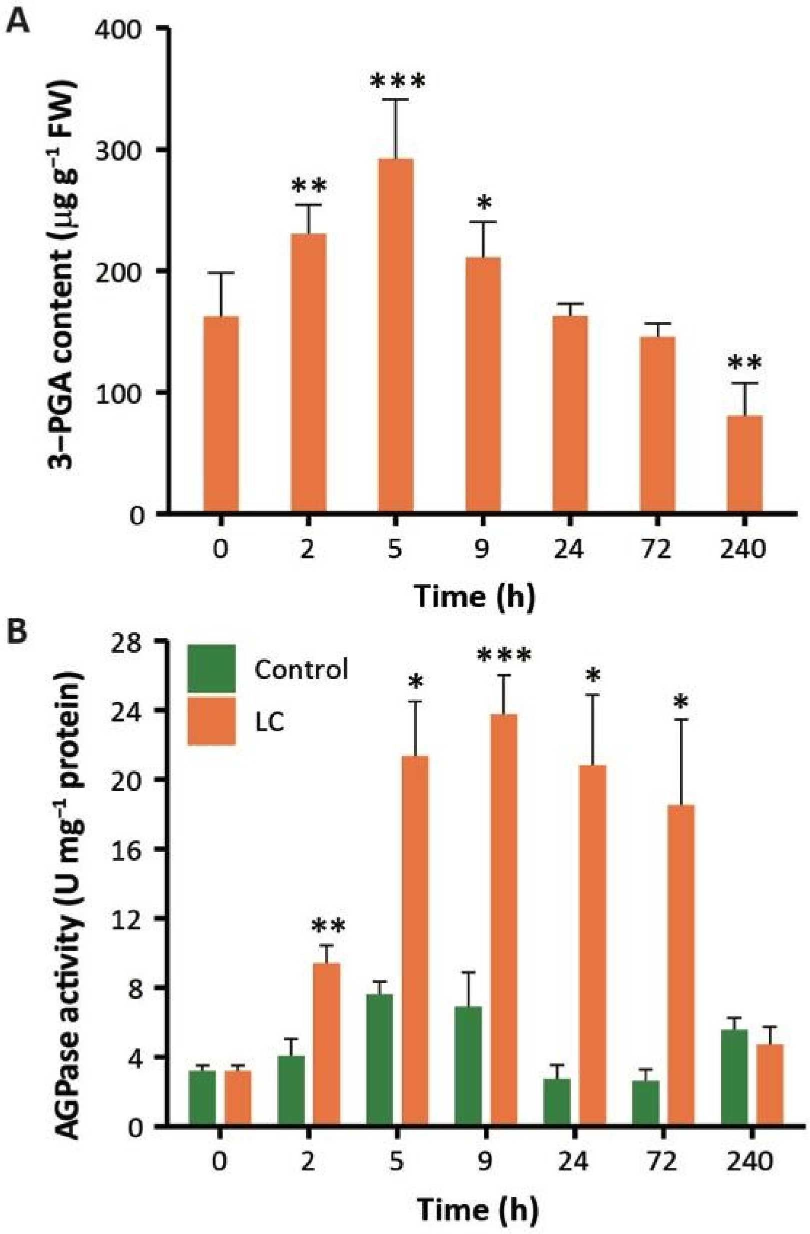


**Fig. S10.**

Changes in 3-PGA content (**A**) and AGPase activity (**B**) in *Landoltia punctata* under LC treatment.

LC, cultivated under conditions of nutrient limitation and elevated $CO_2$ level (2500±100 ppm). Control, cultivated in 1/5 Hoagland medium. 3-PGA, 3-phosphoglycerate. Error bars represent the SDs measured from three independent cultures. Asterisk indicates statistically significant difference between treatment group and 0 h (Student's *t*-test). *, P<0.05; **, P<0.01; ***, P<0.001.

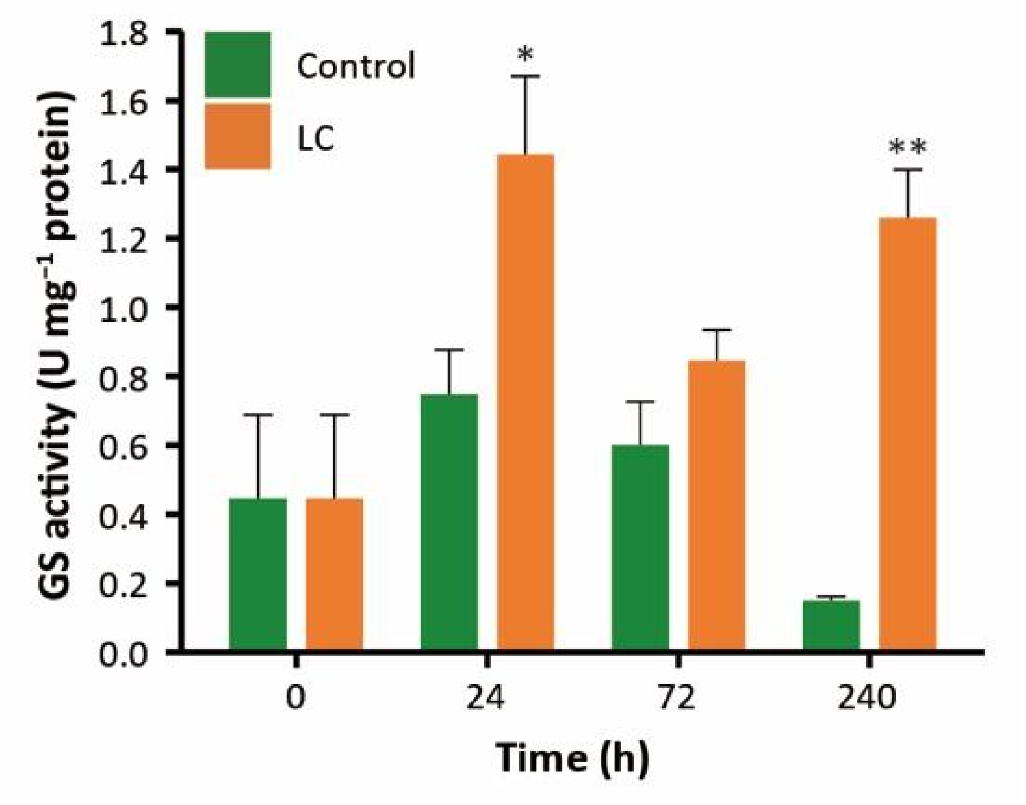


**Fig. S11.**

Activity of GS in *Landoltia punctata*.

LC, cultivated under conditions of nutrient limitation and elevated $CO_2$ level (2500 ± 100 ppm). Control, cultivated in 1/5 Hoagland medium. GS, glutamine synthase. Error bars represent the standard deviation measured from three independent cultures. Asterisks indicate statistically significant difference comparing data from each treatment group with control in the same assay conditions (Student's *t*-test). *, P<0.05; **, P<0.01.

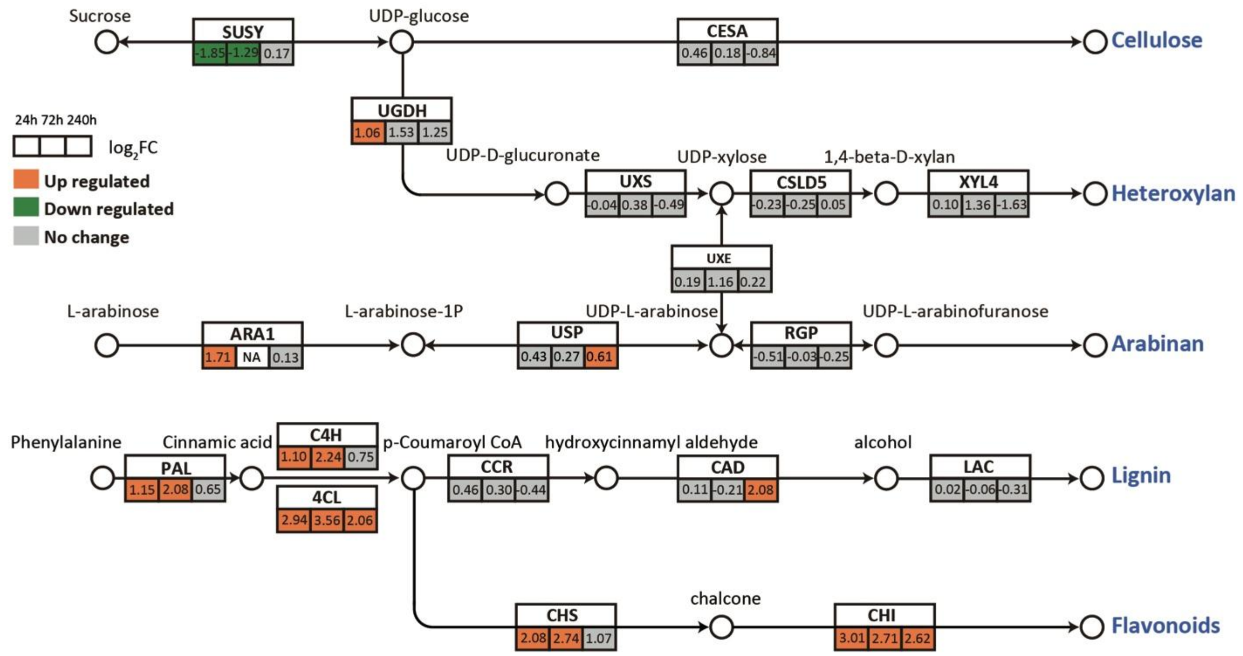


**Fig. S12.**

Expression of key genes involved in biosynthesis of cellulose, hemicellulose, lignin, and flavonoids.

Numbers in boxs are $log_2$(fold change) values at 24 h, 72 h and 240 h after nutrient limitation and elevated $CO_2$ level compared with those at 0 h. Orange or green boxes indicate up- or down-regulated DEGs, respectively. Details are provided in Data S7 and Data S9.

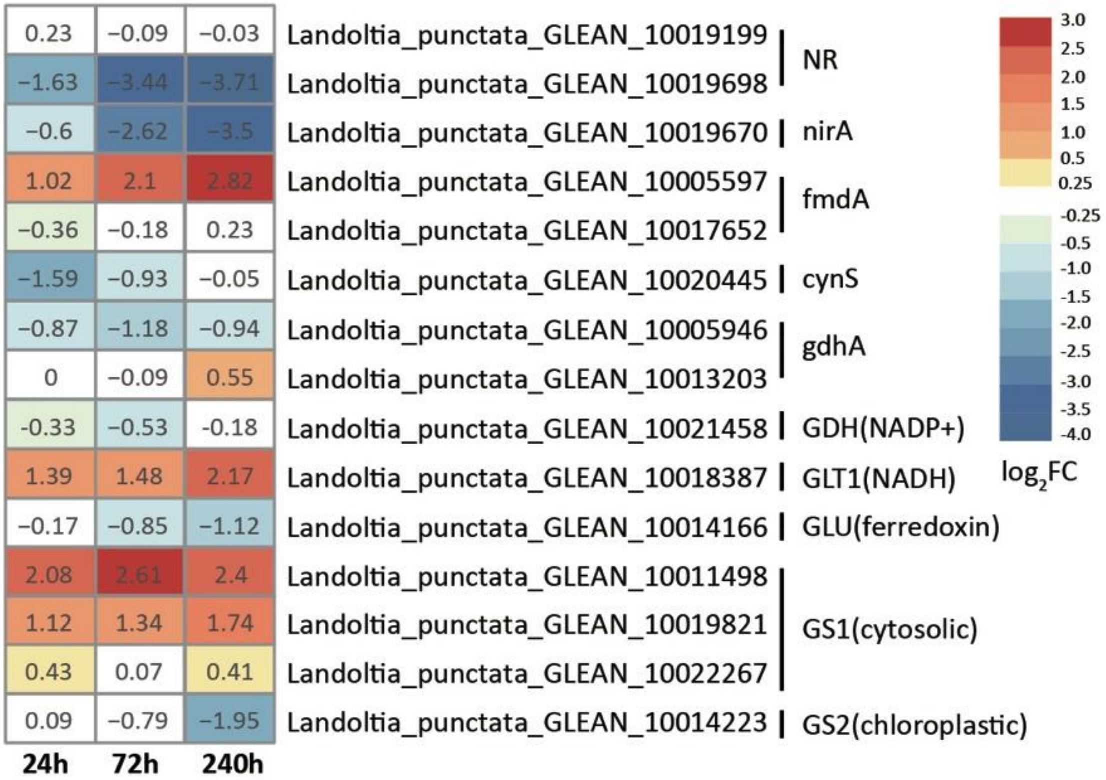


**Fig. S13.**

Expression of genes involved in nitrogen assimilation of *Landoltia punctata*.

Numbers in the boxes are $log_2FC$ values at 24 h, 72 h and 240 h after nutrient limitation and elevated $CO_2$ level compared with expression values (FPKM) with those at 0 h. Details provided in Data S12a.

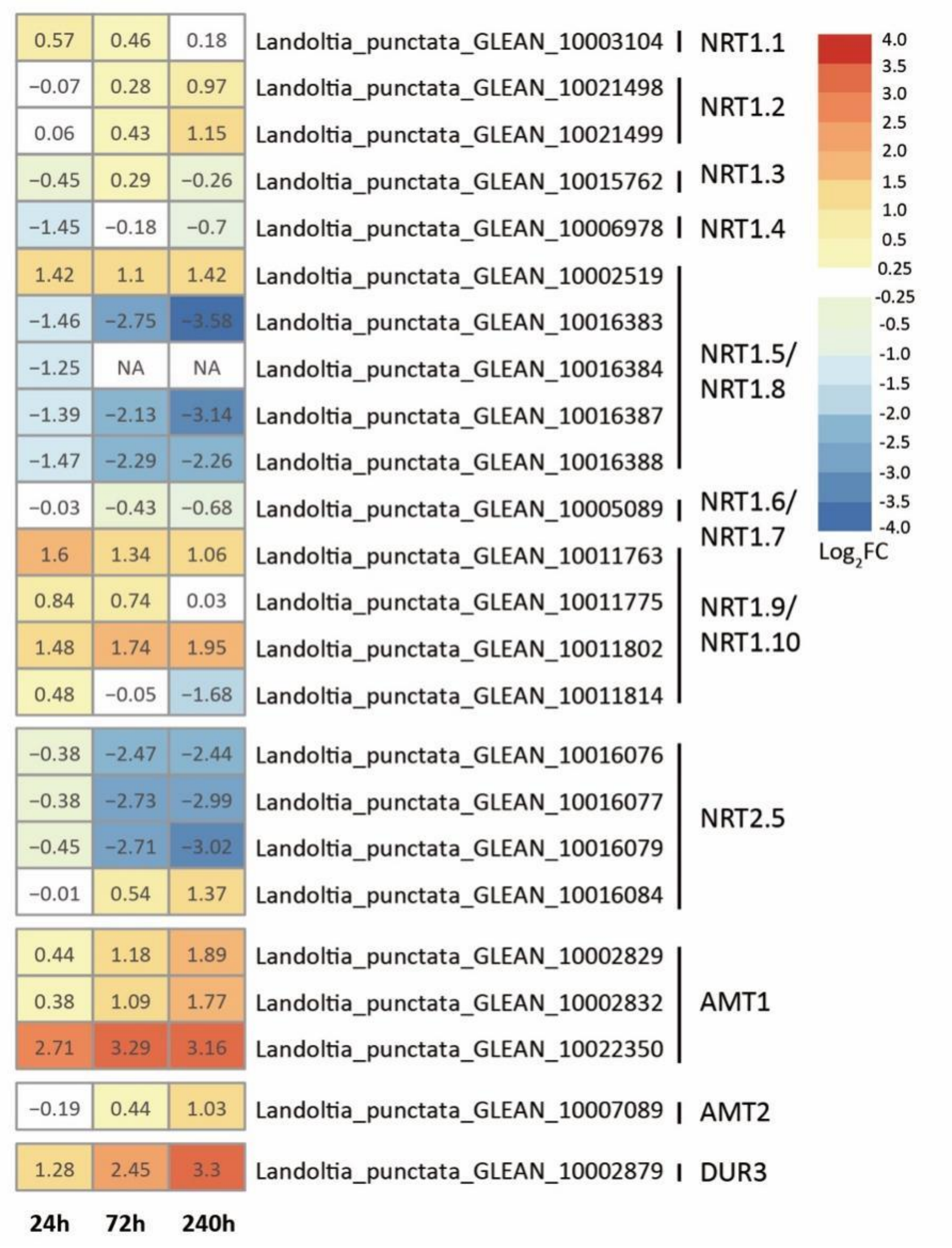


**Fig. S14.**

Expression of genes involved in nitrogen transport of *Landoltia punctata*.

Numbers in the boxes are $\log_2$FC values at 24 h, 72 h and 240 h after nutrient limitation and elevated $CO_2$ level compared with expression values (FPKM) with those at 0 h. Details provided in Data S12b.

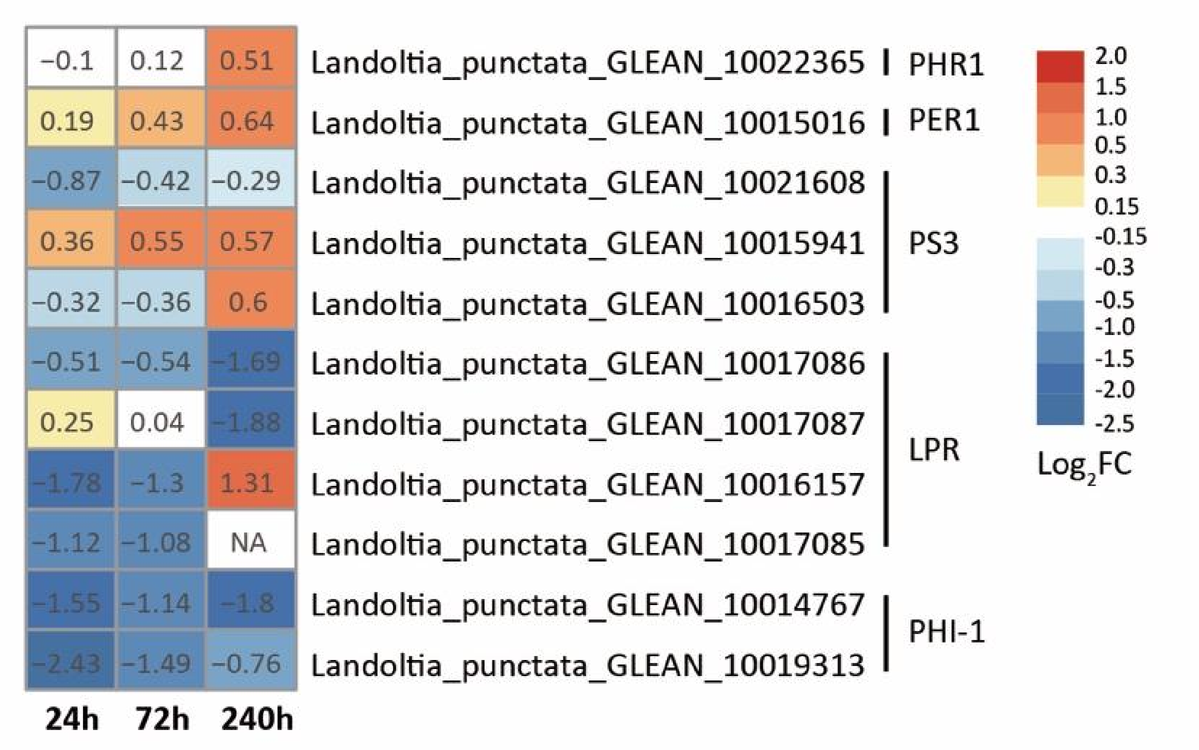


**Fig. S15.**

Expression of genes involved in low phosphate response of *Landoltia punctata*.

Numbers in the boxes are $log_2FC$ values at 24 h, 72 h and 240 h after nutrient limitation and elevated $CO_2$ level compared with expression values (FPKM) with those at 0 h. Details provided in Data S14a.

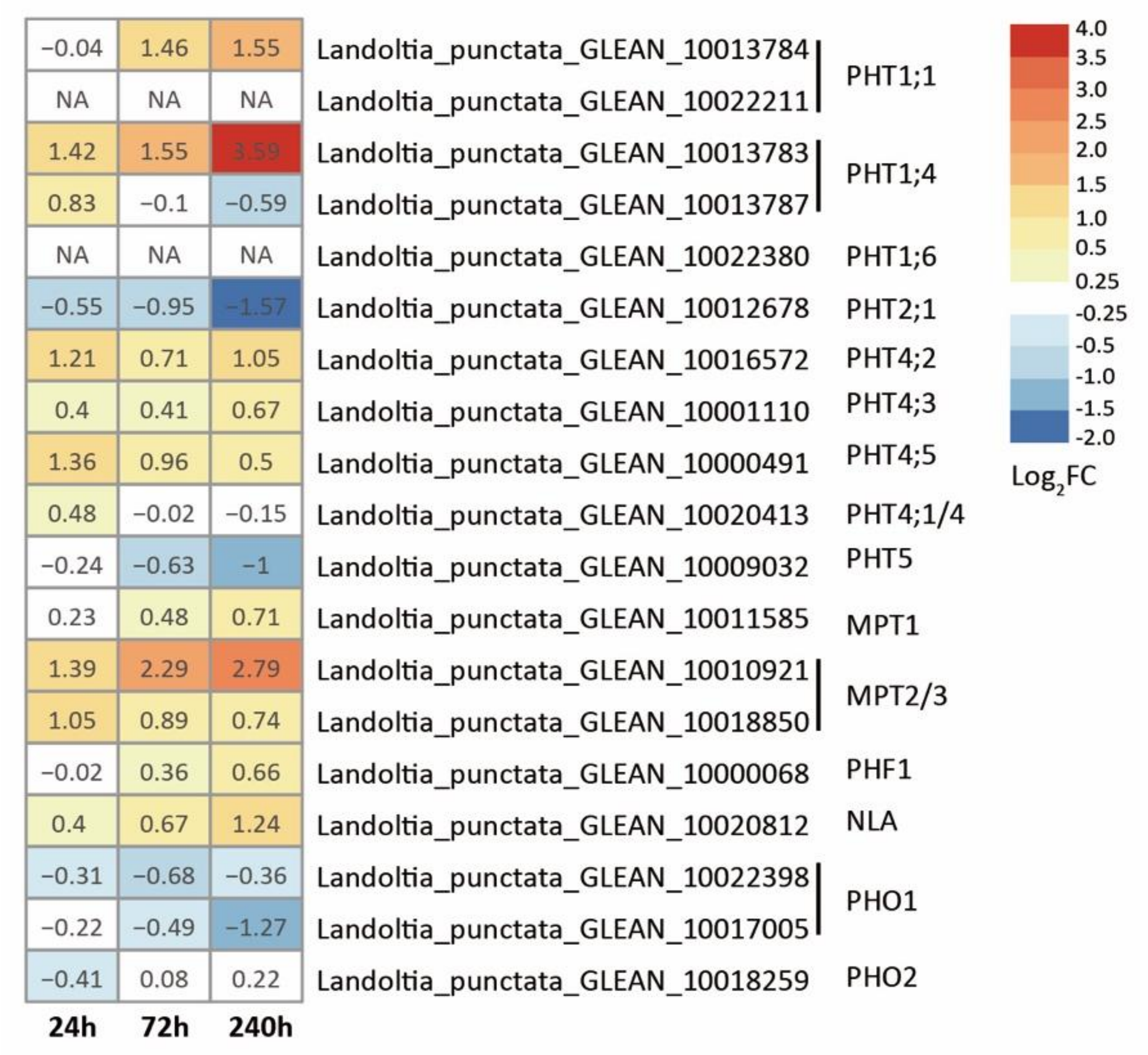


**Fig. S16.**

Expression of genes involved in phosphate transport of *Landoltia punctata*.

Numbers in the boxes are $\log_2$FC values at 24 h, 72 h and 240 h after nutrient limitation and elevated $CO_2$ level compared with expression values (FPKM) with those at 0 h. Details s provided in Data S14b.

A

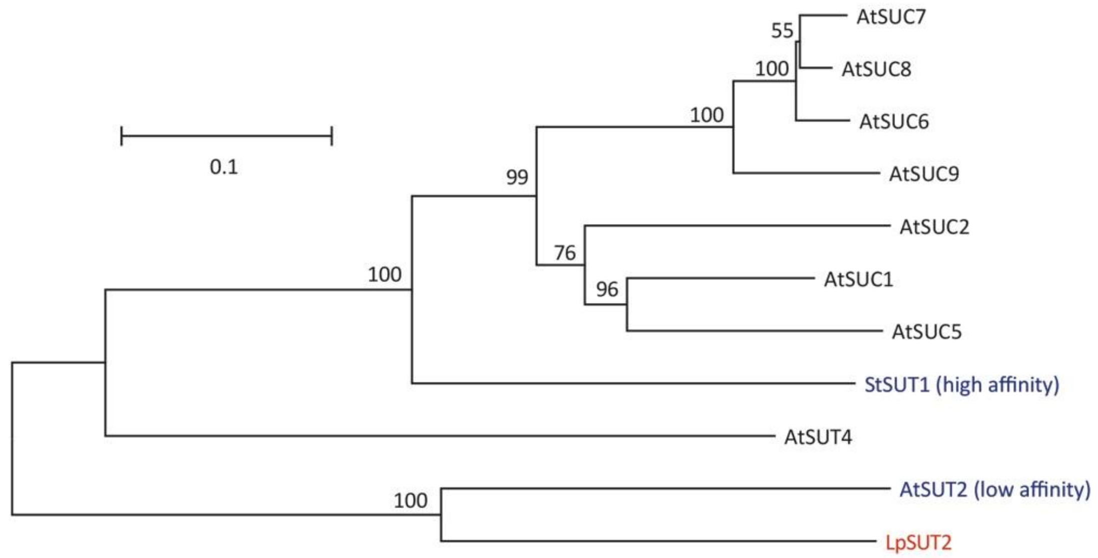


B

```
                                           N-terminal region
StSUT1    1 M------------------ENGTKR-----------EGLGKLTVSSSLQVEQPLAPSKLWKIIVVASIAAGVQFGWALQ 50
AtSUT2    1 MsdsvsisVPYRNLRKEIELETVTKHRQNESgSSSFSESASPSNHSDSADGESVSKNCSLVTLVLSCTVAAGVQFGWALQ 80
LpSUT     1 M-------VSVQDLDTNSEADAGTSRKSNDL-PISLPSSSSPSSAYSSLGMVTEASSSSLKTLILVCMVAAGVQFGWALQ 72

StSUT1   51 LSLLTPYVQLLGIPHKFASFIWLCGPISGMIVQPVVGYYSDNCSSRFGRRRPFIAAGAALVMIAVFLIGFAADLGHASGD 130
AtSUT2   81 LSLLTPYIQTLGISHAFSSFIWLCGPITGLVVQPFVGIWSDKCTSKYGRRRPFILVGSFMISIAVIIIGFSADIGYLLGD 160
LpSUT    73 LSLLTPYIQTLGIEHAFSSFIWLCGPITGLIVQPCVGIWSDNCHSKYGRRRPFIFVGSLLICCAVTIIGFSADLGYMLGD 152

StSUT1  131 TLG-----KGFKPRAIAVFVVGFWILDVANNMLQGPCRALLADLSGgkSGRMRTANAFFSFFMAVGNILGYAAGSYSHLF 205
AtSUT2  161 SKEHCSTFKGTRTRAAVVFIIGFWLLDLANNTVQGPARALLADLSG--PDQRNTANAVFCLWMAIGNILGFSAGASGKWQ 238
LpSUT   153 TKEHCRDYKGPRWRAAAVFIIGFWMLDLANNTVQGPARALLADLSG--HNQQSVANAIFCSWMAVGNILGFSSGSSGQWE 230

StSUT1  206 KVFPFSKTKACDMYCANLKSCFFIAIFLLLSLTTIALTLVRENELPEKDEQEID-------------------------- 259
AtSUT2  239 EWFPFLTSRACCAACGNLKAAFLLAVVFLTICTLVTIYFAKEIPFTSNKPTRIQ-DSAPLLddlqskglehsKLNNGTAN 317
LpSUT   231 RWFPFLRTRACCEACGNLKAAFLVAVAFLMFCTVVTLYFAVEVPLVKVNPENTSsDSAPLL-----------KVDLKDVN 299
                           Central loop
StSUT1  260 -EKLAGAGKSKVPFFG---------------------EIFGALKELPRPMWILLLVTCLNWIAWFPFFLYDTDWMAKEVF 317
AtSUT2  318 GIKYERVERDTDEQFGnSENE-HQDETYVDGPGSVLVNLLTSLRHLPPAMHSVLIVMALTWLSWFPFFLFDTDWMGREVY 396
LpSUT   300 GLQIASSEK--------SEKEvNQLAPFDNSPSSVLVNLLTSLRHLPPGMTSVLLVMSLTWLSWFPFFLFDTDWMGREVY 371

StSUT1  318 -GGQVGDA---RLYDLGVRAGAMGLLLQSVVLGFMSLGVEFLGKKIGgAKRLWGILNFVLAICLAMTILVTKMaEKSRQH 393
AtSUT2  397 HGDPTGDSLHMELYDQGVREGALGLLLNSVVLGISSFLIEPMCQRMG-ARVVWALSNFTVFACMAGTAVISLM-SLSDDK 474
LpSUT   372 HGNPTGDVDEVNLYHHGVRAGAFGLLLNSVVLMVSSFLIEPMCRRLG-PRLVWALSNFVVCFCMTATAVISLL-SLAEYS 449

StSUT1  394 DPAGTLMGPTPGVKIGALLLFAALGIPLAATFSIPFALASIFSSNRGSGQGLSLGVLNLAIVVPQMLVSLVGGPWDDLFG 473
AtSUT2  475 NGIEYIMRGNETTRTAAVIVFALLGFPLAITYSVPFSVTAEVTADSGGGQGLAIGVLNLAIVIPQMIVSLGAGPWDQLFG 554
LpSUT   450 ESVQHMIQGNGGHKVAALVLFALLGFPLAITYSVPYSMTAELTAGSGGGQGLATGVLNLAIVIPQMIVSLGAGPWDALFG 529

StSUT1  474 GGNLPGFVVGAVAAAASAVLALTMLPSPPADAKPAVAMGLSik 516
AtSUT2  555 GGNLPAFVLASVAAFAAGVIALQRLPTLSSSFKSTGFH-IG-- 594
LpSUT   530 GGNMPAFVLAAAFSLAAGIVSVLKLPGLVSGSHMSGFHGFG-- 570
```

C

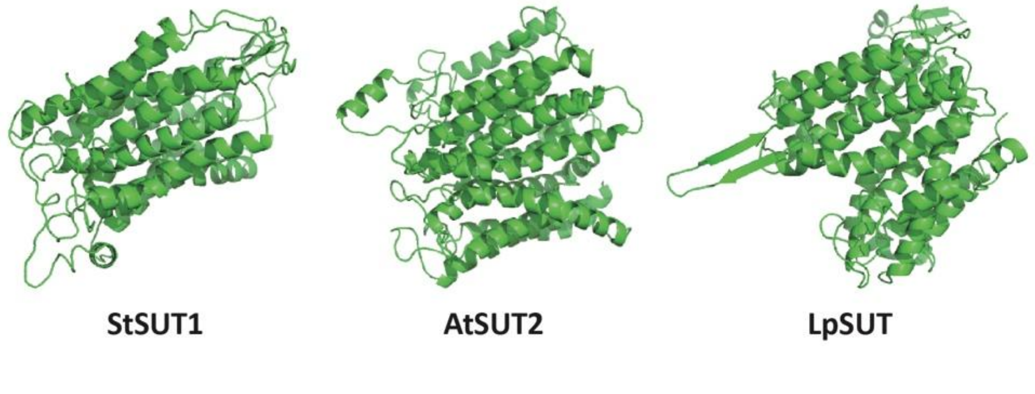

**Fig. S17.**

SUT phylogenetic tree and sequence alignment among *Landoltia punctata* (Lp), *Arabidopsis thaliana* (At), and *Solanum tuberosum* (St).

**A**, SUT phylogenetic tree. Bar, substitution/site, 0.1.

**B**, Sequence alignment of SUT proteins among *Landoltia punctata* (*LpSUT*), *Arabidopsis thaliana* (*AtSUT2*, low-affinity sucrose transporter), and *Solanum tuberosum* (*StSUT1*, high-affinity sucrose transporter). Cytoplasmic extended domain at N-terminal region (first black line region) and extended domain at central loop region (second black line region) are marked in red boxes.

**C**, Predicted 3D structure of StSUT1, AtSUT2, and LpSUT.

**A**

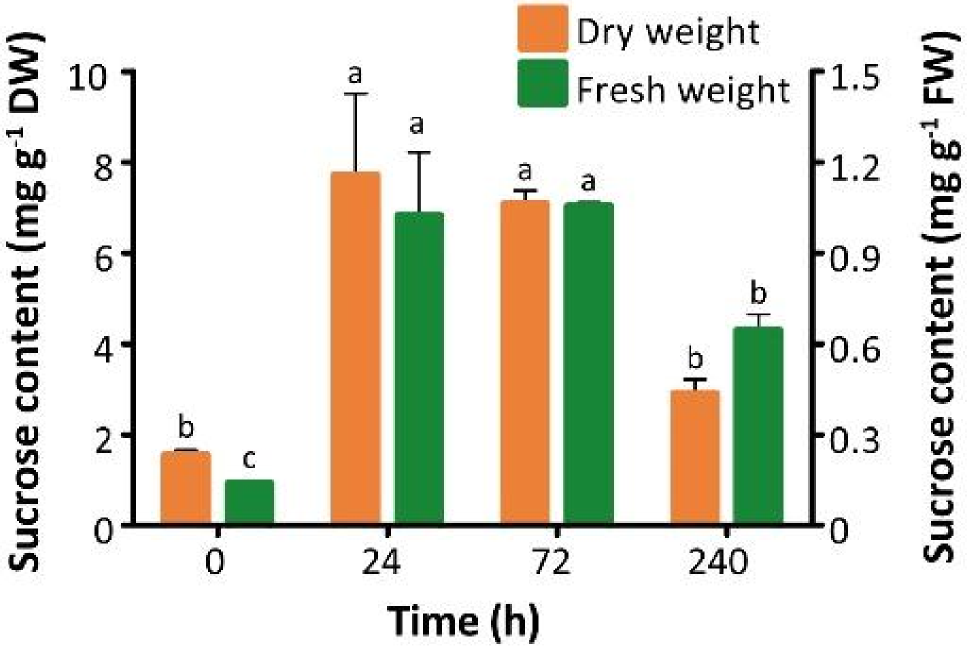


**B**

| Species | Tissue | Sucrose content | |
|---|---|---|---|
| | | mg $g^{-1}$ DW | mg $g^{-1}$ FW |
| Rice | Leaf shade | 45 | -- |
| maize | Leaf shade | 20 | -- |
| *Arabidopsis thaliana* | Rosette leaf | -- | 2.6-2.7 |
| *Hevea brasiliensis* | Leaf | 38-65 | -- |
| *Landoltia punctata* | Frond | 1.5-7.7 | 0.1-1.1 |

**Fig. S18.**

Sucrose contents in *Landoltia punctata* under LC treatment.

**A**, Sucrose content in *Landoltia punctata*. Letters indicate significant differences among time points. Dry weights and Fresh weights are tested by one way ANOVA following Tukey-Kramer test ($p<0.05$).

**B**, Comparison of sucrose content in *Landoltia punctata* (determined in this study), rice (*3*), maize (*4*), *Arabidopsis thaliana (5),* and *Hevea brasiliensis (6).*

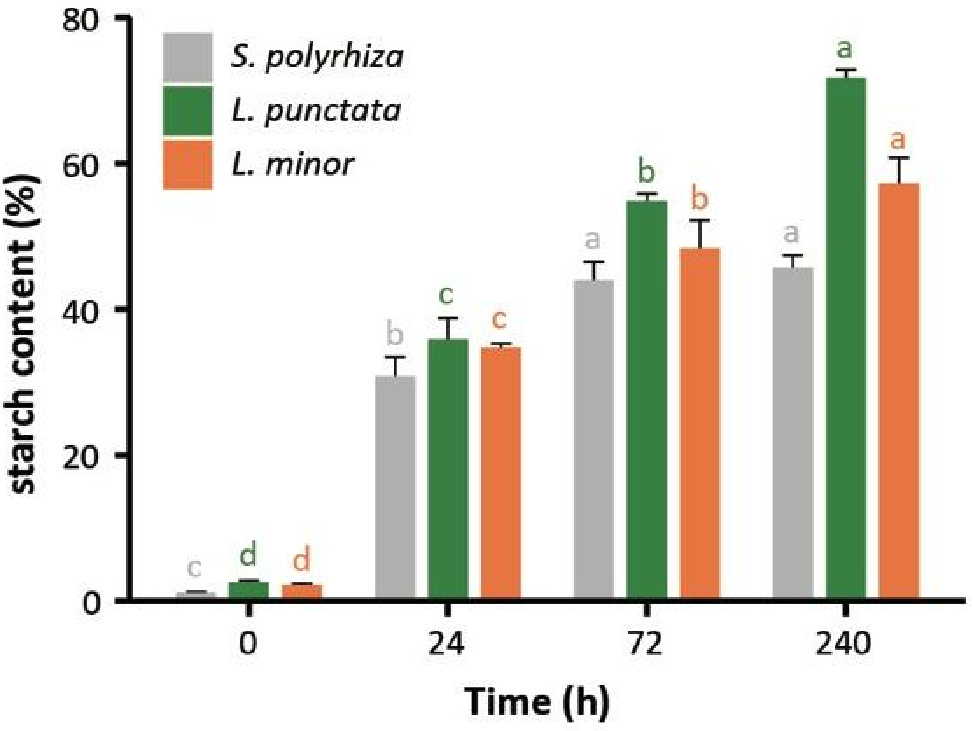


**Fig. S19.**

Accumulation of starch in *Spirodela polyrhiza*, *Landoltia punctata*, and *Lemna minor* under LC treatment.

Error bars show standard deviations measured from three independent cultures. Letters indicate significant differences determined by one way ANOVA following Tukey-Kramer test ($p<0.05$).

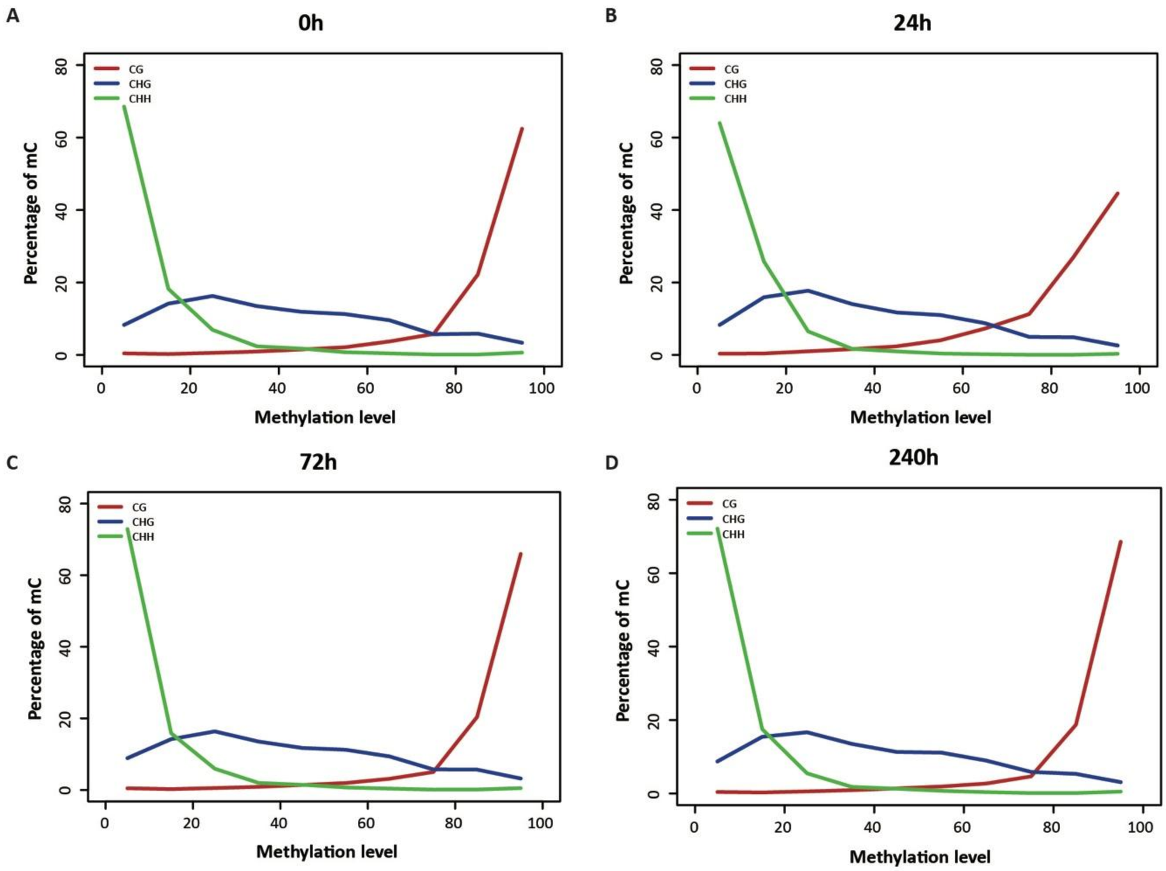


**Fig. S20.**

Methylation level distribution of methylated cytosine (mC).

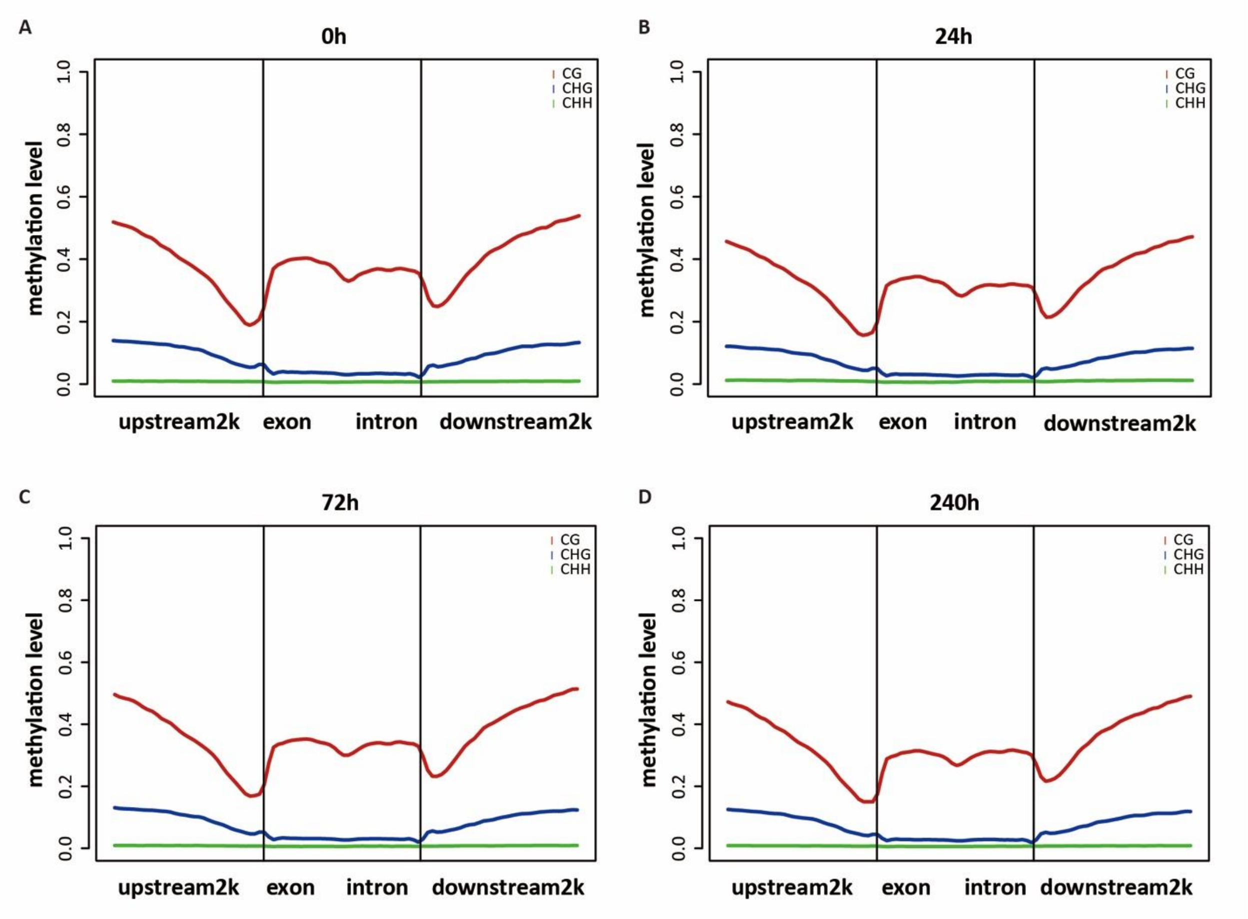


**Fig. S21.**

The methylation level of different regions of genome. Upstream2k/downstream2k, the 2 kb-upstream or 2 kb-downstream regions of genes.

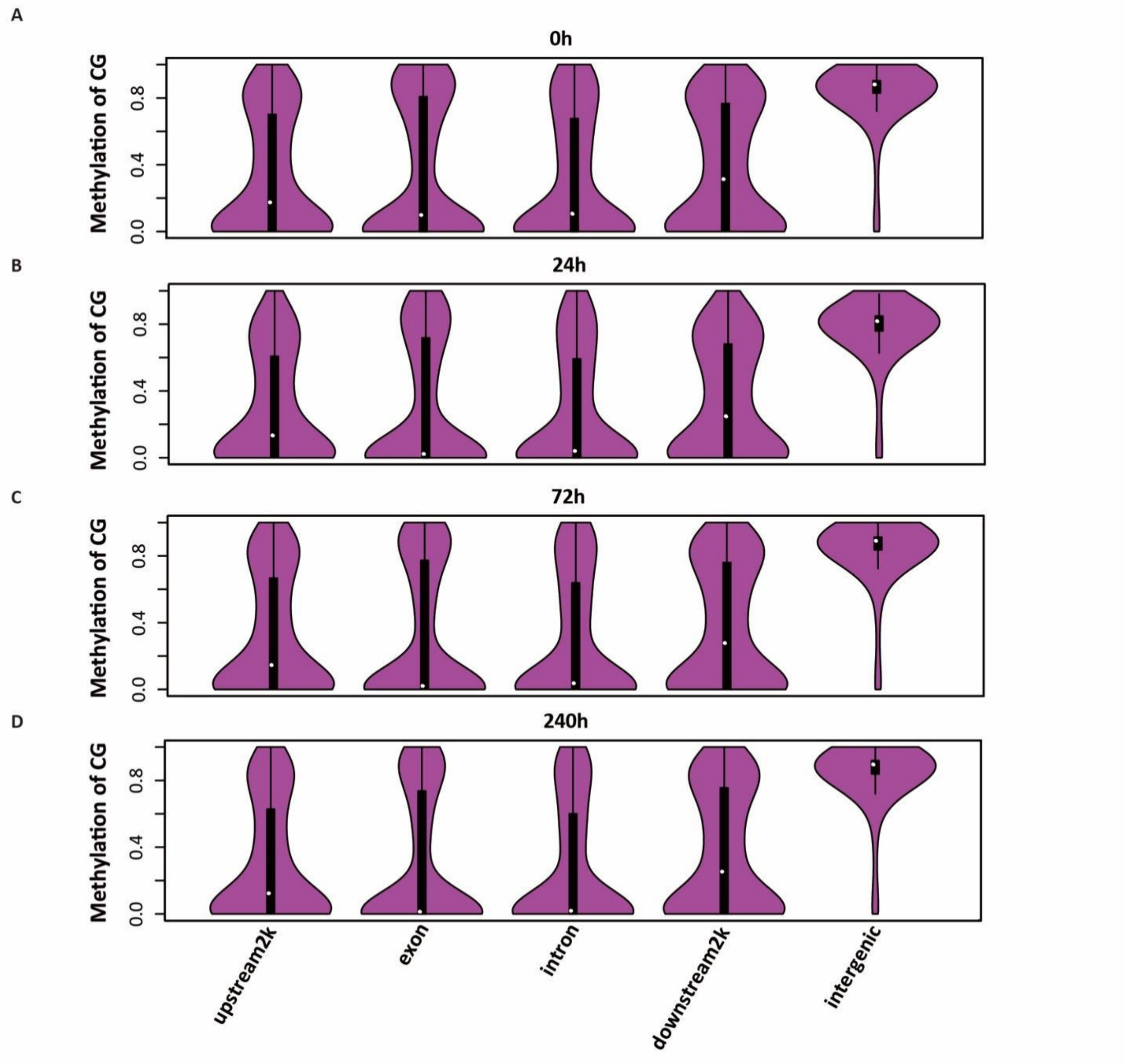


**Fig. S22.**

Methylation level of CG in different regions of genome. Upstream2k/downstream2k, the 2 kb-upstream or 2 kb-downstream regions of genes.

**Table S1.**

Treatments for starch production by *Landoltia punctata*.

| Treatment | $CO_2$ concentration (ppm) | Media |
|---|---|---|
| Control | 390±50[2] | 1/5 Hoagland |
| C[1] | 2500±100 | 1/5 Hoagland |
| L[1] | 390±50[2] | Deionized water |
| LC[1] | 2500±100 | Deionized water |

[1] C: elevated $CO_2$ level; L, nutrient limitation; LC, nutrient limitation and elevated $CO_2$ level.

[2] The $CO_2$ concentration in atmosphere is 390±50 ppm.

**Table S2.**

RNA-Sequencing data.

| Sample ID | Raw bases (Gb) | Clean bases (Gb) | Read length (bp) | Base Q20 (%) | GC content (%) |
|---|---|---|---|---|---|
| LC0h_1 | 7.2 | 6.9 | 150×2 | 97.8 | 56.4 |
| LC0h_2 | 7.2 | 6.9 | 150×2 | 97.8 | 56.5 |
| LC0h_3 | 7.2 | 6.9 | 150×2 | 97.8 | 55.3 |
| LC24h_1 | 7.2 | 7.0 | 150×2 | 98.0 | 57.1 |
| LC24h_2 | 7.2 | 7.0 | 150×2 | 97.6 | 56.7 |
| LC24h_3 | 7.2 | 7.0 | 150×2 | 97.6 | 56.7 |
| LC72h_1 | 7.2 | 7.0 | 150×2 | 98.0 | 56.9 |
| LC72h_2 | 7.2 | 7.0 | 150×2 | 97.2 | 56.7 |
| LC72h_3 | 7.2 | 6.9 | 150×2 | 97.7 | 56.5 |
| LC240h_1 | 7.0 | 6.9 | 150×2 | 98.0 | 56.1 |
| LC240h_2 | 7.2 | 7.0 | 150×2 | 98.0 | 56.2 |
| LC240h_3 | 7.2 | 6.9 | 150×2 | 98.0 | 55.8 |

**Table S3.**

Primers for the selected DEGs for qRT-PCR.

F: Forward primer; R: Reverse primer.

| Gene ID | Primers (5'→3') | Products length (bp) |
|---|---|---|
| Landoltia_punctata_GLEAN_10003767 | F: GGATCGGGCTGGTCGTTAC<br>R: GAATTCCCCCTTCTTGCTG | 96 |
| Landoltia_punctata_GLEAN_10001924 | F: CATTGGTGCTGTTTTTTGAAGGG<br>R: TTCATCATTTGCAGCTACGATCC | 97 |
| Landoltia_punctata_GLEAN_10003150 | F: GAACTCCCACTCGCTGTTGTCT<br>R: CCTCTCGGTACTCTTCTCCCCT | 116 |
| Landoltia_punctata_GLEAN_10015586 | F: CGCCGTCCTTCAAACCAA<br>R: CCAATACGCCGGGCCATA | 148 |
| Landoltia_punctata_GLEAN_10000800 | F: TTTTTCAGGAGAGAGGTGTCCA<br>R: GGTTTGAGCAGTTTGATGGC | 128 |
| Landoltia_punctata_GLEAN_10004280 | F: TTCGCGACTCCTGTTTCGTT<br>R: ATACCTGGCGTGCGTGTTCA | 106 |
| Landoltia_punctata_GLEAN_10011855 | F: GCTATCTGCCCAAGGTCTGC<br>R: GGGGGTTCTCCACTTCCACT | 126 |
| Landoltia_punctata_GLEAN_10019540 | F: CAGCCGTTCACGTCTTTGGT<br>R: TGCCGTCCTCGTCCTTCTTA | 114 |
| Landoltia_punctata_GLEAN_10019599 | F: AGCAAGAGCCCCGTCACC<br>R: CCGGCGCAACTTCCAAT | 113 |
| Landoltia_punctata_GLEAN_10000438 | F: CCTCGTTTTGGGTTGTAATATTC<br>R: GGCAAGGGACCTCCTTTGGC | 104 |
| Landoltia_punctata_GLEAN_10012804 | F: ATGCTTTGACAGAGCGTGTTG<br>R: GTTGCTTAGGCTGCCAGTG | 121 |
| Landoltia_punctata_GLEAN_10000828 | F: TTTCGGGCTGTTCGGGTTG<br>R: GGCGATGATGGCGTTCTTCT | 94 |

| | | |
|---|---|---|
| Landoltia_punctata_GLEAN_10018179 | F: TGCCGTCCTCGCCTTACTG<br>R: CCGCAGCCGTTCACATCTT | 117 |
| Landoltia_punctata_GLEAN_10004572 | F: GGGAAAAAAGTCGAGCATACC<br>R: TCGTTGCCCACAGTGAAAATA | 133 |
| Landoltia_punctata_GLEAN_10010520 | F: TAGCGGAGTAAAATCGGTGACG<br>R: GGGGAAATGAAAAATGGAGAAA | 116 |
| Landoltia_punctata_GLEAN_10019316 | F: ACCCAACTCCCTGTCATTTTTCT<br>R: CCTGCTTTATCCGGTTCCATC | 109 |
| Landoltia_punctata_GLEAN_10011880 | F: TCTCTCGTGCTTTCTTCGCTC<br>R: GCACACCTTCTTCCACTCCATC | 90 |
| Landoltia_punctata_GLEAN_10012751 | F: GTCGCTGTGATGCTGGTTTTC<br>R: TTCCCGTGTACATGACGTTGC | 133 |
| Landoltia_punctata_GLEAN_10014601 | F: CTCGGCGAGGAACACGCT<br>R: TCCCAGATGCCCAAACGG | 140 |
| Landoltia_punctata_GLEAN_10018050 | F: TACGGGGAGGGAGAACCC<br>R: CGCCGACGACGACAACA | 133 |
| *Actin* (Internal control) | F: TGATGGTTGGAATGGGACAG<br>R: TTGGTCACAACGCCATGCT | 106 |

**Table S4.**

Primers for quantification of the key genes expression involved in $CO_2$ fixation, carbon concentration, and starch synthesis by qRT-PCR.

F: Forward primer; R: Reverse primer.

| Gene name | Primers (5'→3') | Products length (bp) |
|---|---|---|
| *Actin* (Internal control) | F: TGATGGTTGGAATGGGACAG<br>R: TTGGTCACAACGCCATGCT | 106 |
| *AGPase* | F: ATCACGCATCCTTCACCAATC<br>R: TCATCTCCAATCTACACTCAACCT | 95 |
| *SSS* | F: TATGGCACGGAAGAGTTGAAG<br>R: TCTCTCTGGCGAAGGAACTCA | 151 |
| *Rubisco* | F: GGTGGAGGAGGTCAAGAAGG<br>R: GCTTGGCTGCAATGAAACTG | 101 |
| *PEPC* | F: GAGATGAGAGCGGGGATGAG<br>R: TGAGAGGAGCGTTGTAGGGG | 130 |
| *GBSS* | F: GGACCCCAACGCTCTTCTCTT<br>R: CGACCATCTGGATGTTCTCGC | 113 |
| *UGPase* | F: TTGAAGGTATCTGGGGATGTGTG<br>R: GAGTTGACTTGTTCTCGAGGGTG | 127 |

**Table S5.**

WGBS data of *Landoltia punctata* under LC treatment.

Samples were withdrawn at time points of 0 h, 24 h, 72 h, and 240 h, which were marked as LC_0 h, LC_24 h, LC_72 h, and LC_240 h, respectively. RL, read length; $R_a$, align rate; $R_{nd}$, nonDup rate; $R_c$, conversion rate.

| Samples | Raw reads | Clean reads | RL (bp) | Q20 (%) | Align reads | $R_a$ (%) | nonDup reads | $R_{nd}$ (%) | $R_c$ (%) |
|---|---|---|---|---|---|---|---|---|---|
| LC_0 h | 165,497,180 | 151,982,152 | 150×2 | 99.7 | 131,841,668 | 86.8 | 119,789,431 | 90.9 | 96.9 |
| LC_24 h | 178,456,582 | 165,815,374 | 150×2 | 99.6 | 128,862,610 | 77.7 | 120,079,561 | 93.2 | 96.7 |
| LC_72 h | 195,165,098 | 179,820,248 | 150×2 | 99.7 | 158,551,840 | 88.2 | 143,195,052 | 90.3 | 94.5 |
| LC_240 h | 204,367,354 | 189,216,976 | 150×2 | 99.7 | 166,590,668 | 88.0 | 150,834,753 | 90.5 | 95.5 |

**Table S6.**

The global methylation level of *Landoltia punctata* under LC treatment.

Samples were withdrawn at time points of 0 h, 24 h, 72 h, and 240 h, which were marked as LC_0 h, LC_24 h, LC_72 h, and LC_240 h, respectively.

| Samples | C (%) | CG (%) | CHG (%) | CHH (%) |
|---|---|---|---|---|
| LC_0 h | 12.9 | 76.7 | 22.0 | 1.4 |
| LC_24 h | 11.2 | 70.6 | 19.7 | 1.8 |
| LC_72 h | 13.0 | 77.4 | 21.9 | 1.3 |
| LC_240 h | 12.9 | 78.1 | 22.2 | 1.3 |

**Table S7.**

Starch production ability of *Landoltia punctata* in pilot scale.

Fresh duckweed of approximately 4.0 kg was inoculated into 3.1×4.5×0.4 $m^3$ (W×L×D) tanks filled with tap water. Duckweed was cultivated at 25ºC with sunlight in daytime and fluorescent lamp at night. $CO_2$ was aerated to a concentration of 2500±100 ppm.

| Batch | 1 | 2 | 3 | 4 | Mean |
|---|---|---|---|---|---|
| Cultivation time (d) | 4 | 4 | 4 | 4 | - |
| Starch content (% DW) | 41.25 | 49.34 | 45.48 | 47.66 | 45.93±3.50 |
| Growth rate (g $m^{-2}$ $d^{-1}$) | 14.31 | 14.63 | 16.98 | 17.92 | 15.96±1.77 |
| Starch productivity (g $m^{-2}$ $d^{-1}$) | 8.55 | 9.24 | 10.89 | 11.48 | 10.04±1.37 |
| Moisture (%) | 84.35 | 81.12 | 83.47 | 83.14 | 83.02±1.36 |

**Table S8.**

Links for genomes used in this research.

| Species | Links |
|---|---|
| *Klebsormidium flaccidum* | http://www.plantmorphogenesis.bio.titech.ac.jp/~algae_genome_project/klebsormidium/kf_download.htm |
| *Spirodela polyrhiza* | ftp://ftp.ncbi.nlm.nih.gov/genomes/all/GCA/001/981/405/GCA_001981405.1_ASM198140v1/GCA_001981405.1_ASM198140v1_genomic.gff.gz |
| *Landoltia punctata* | https://www.ncbi.nlm.nih.gov/bioproject/PRJNA546087 |
| *Zostera marina* | ftp://ftp.ncbi.nlm.nih.gov/genomes/all/GCA/001/185/155/GCA_001185155.1_Zosma_marina.v.2.1/GCA_001185155.1_Zosma_marina.v.2.1_genomic.gff.gz |
| *Lemna minor* | https://genomevolution.org/GenomeInfo.pl?gid=27419 |
| *Arabidopsis thaliana* | ftp://ftp.ncbi.nlm.nih.gov/genomes/all/GCF/000/001/735/GCF_000001735.3_TAIR10/GCF_000001735.3_TAIR10_genomic.gff.gz |
| *Oryza sativa* | ftp://ftp.ncbi.nlm.nih.gov/genomes/all/GCF/000/005/425/GCF_000005425.2_Build_4.0/GCF_000005425.2_Build_4.0_genomic.gff.gz |
| *Zea mays ssp. mays* | ftp://ftp.ncbi.nlm.nih.gov/genomes/all/GCF/000/005/005/GCF_000005005.2_B73_RefGen_v4/GCF_000005005.2_B73_RefGen_v4_genomic.gff.gz |
